# Carbohydrate utilization regulator reveals a noncanonical mechanism of nutrient differentiation

**DOI:** 10.1101/2025.04.10.648202

**Authors:** Brandon Reyes-Chavez, Joshua D. Kerkaert, Lori B. Huberman

## Abstract

Cells must sense and respond to nutrients to survive. To efficiently grow in mixed carbon environments, microbes repress genes necessary to utilize carbon sources that require substantial resources to catabolize when a simpler carbon source, such as glucose, is present. This process is known as carbon catabolite repression. Canonically, in fungi, nutrient sensing transcriptional networks are composed of carbon source-specific transcription factors that activate carbon source utilization genes and carbon catabolite repression regulators, which broadly repress all nonpreferred carbon source utilization genes when a preferred carbohydrate is present. In contrast to this model, we identified a transcription factor (Cbr1) in the basidomycete yeast *Rhodotorula* (*Rhodosporidium*) *toruloides* that specifically inhibits glucose-mediated repression of glucose-glucose disaccharide utilization, presenting a mechanism of tailored carbon catabolite repression regulation that combats a negative feedback loop formed when glucose is released during disaccharide utilization. Cbr1 is also required for cellobiose, carboxylic acid, and fucose utilization. Using transcriptomic and molecular analyses, we demonstrated that catabolism of these carbon sources is not metabolically linked, but genes necessary for their utilization are coactivated by Cbr1 in response to each of the carbon sources. This coactivation suggests *R. toruloides* may encounter these carbon sources together, potentially during complex interactions among microbes in nature. Coregulation of nutrient-specific gene activation and carbon catabolite repression by a transcription factor establishes a previously uncharacterized mechanism for building nutrient sensing transcriptional networks in fungi. Characterizing diverse nutrient sensing regulatory mechanisms is critical for understanding resource acquisition during fungal pathogenesis, where carbon catabolite repression is important for virulence and drug tolerance, and metabolically engineering fungi for green biotechnology.

## INTRODUCTION

Cells must sense and respond to nutrients to successfully grow and divide using the resources available. For fungi, nutrient sensing is critical to establish fungal colonies, compete with other microbes, and establish pathogenic and symbiotic relationships with plants and animals. Most work characterizing genetic pathways regulating carbon source utilization in fungi has been performed in the Ascomycete phylum [1, 2]. In these fungi, a network of carbon source-specific transcription factors activates genes necessary for utilization of available carbon [3]. However, in a mixed carbon environment, it is not necessarily energetically optimal to simultaneously consume all available carbon sources. Thus, when a more preferred carbon source, such as glucose, is available, a process known as carbon catabolite repression broadly downregulates expression of genes required for utilization of all less preferred carbon sources [1, 3–10].

Many devastating human, animal, and plant pathogens fall outside of the Ascomycete phylum [11]. Genetic mechanisms through which these fungi activate genes necessary to utilize nonpreferred carbon sources and repress these genes in the presence of more preferred carbon sources are significantly less well studied. Basidiomycetes diverged from Ascomycetes approximately 400 million years ago and represent over 30% of described fungi [12, 13]. Recent studies in Basidiomycete pathogens have demonstrated that understanding the regulation of carbon utilization in this phylum is critical, since accurately regulating carbon metabolism is not only important for virulence but also changes the efficacy of antifungal drug treatments [14–16].

To address this knowledge gap, we investigated genetic mechanisms regulating carbon utilization in the basidiomycete yeast *Rhodotorula toruloides* (previously *Rhodosporidium toruloides*). Recent studies of Ascomycete yeast species have established these organisms have a range of carbon utilization repertoires, from specialists capable of consuming only a small range of carbohydrates to generalists that have a much broader carbon utilization repertoire [17]. *R. toruloides* is a generalist that can consume most soluble breakdown products of plant biomass [18, 19]. *R. toruloides* can also accumulate up to 70% of its biomass as fatty acids in conditions in which carbon is abundant but other nutrients are limiting and produces useful secondary metabolites, such as carotenoids [20–26]. This makes *R. toruloides* an excellent choice for metabolic engineering [27], where understanding the regulation of carbon metabolism is critical [28, 29].

Cellulose is the most abundant plant biomass component and is composed of glucose chains linked by β-1,4-glycosidic bonds [30]. *R. toruloides* can utilize the disaccharide cellulose building block, cellobiose [31]. We identified a transcription factor, Cbr1, required for cellobiose utilization. Cbr1 is homologous to the cellulose degradation regulator CLR-2/ClrB in Ascomycetes [32], but the role of Cbr1 is significantly expanded in *R. toruloides*, elucidating alternative mechanisms of building carbon utilization regulatory networks in fungi. Unlike CLR-2/ClrB, Cbr1 is also required for tricarboxylic acid (TCA) cycle intermediate and fucose utilization. Cellobiose, fucose, and TCA cycle intermediates are not catabolized through similar pathways, suggesting *R. toruloides* may inhabit an ecological niche in which these carbon sources are frequently found together, potentially through cooperation with or exploitation of saprophytic filamentous fungi.

Further investigation of Cbr1 demonstrated this transcription factor inhibits carbon catabolite repression specifically of glucose-glucose disaccharide utilization genes. This role for Cbr1 contrasts with previously identified regulatory mechanisms of carbon catabolite repression, which regulate utilization of all nonpreferred carbon sources. Taken together, the diverse roles of Cbr1 in *R. toruloides* metabolic regulation present a previously uncharacterized mechanism of carbon catabolite repression and nutrient sensing transcriptional network composition in fungi.

## RESULTS

### Cbr1 is required for wild-type utilization of cellobiose, TCA cycle intermediates, and fucose

*R. toruloides* can utilize cellobiose, the disaccharide breakdown product of the most abundant plant cell wall polysaccharide, cellulose [31]. Several transcription factors regulate cellulose and cellobiose utilization in Ascomycete filamentous fungi, including CLR-1/ClrA, CLR-2/ClrB, and XlnR [1, 32–35]. To investigate genetic mechanisms regulating *R. toruloides* cellobiose utilization, we searched for genes in the *R. toruloides* genome with homology to these transcription factors. No clear homolog of CLR-1/ClrA or XlnR existed in *R. toruloides* (Fig S1). However, we identified a single homolog of CLR-2/ClrB, RTO4_13588 (Fig 1A and Fig S1).

**Fig 1.**
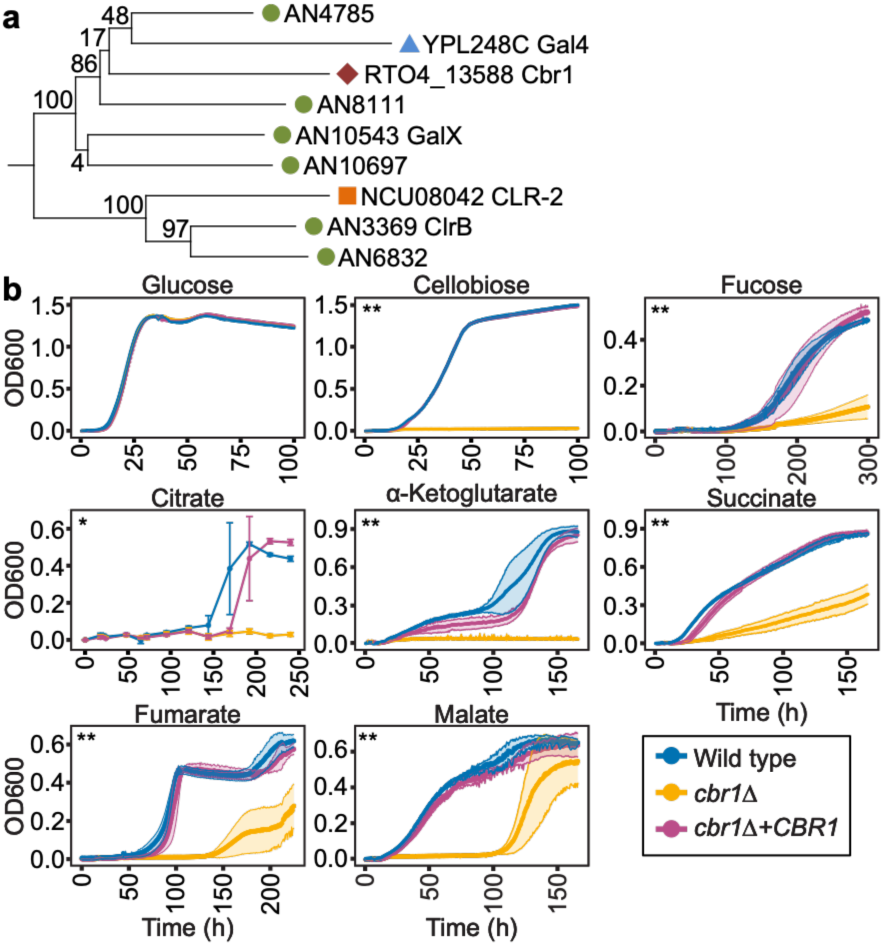
*CBR1* (RTO4_13588) is involved in utilization of cellobiose, TCA cycle intermediates, and fucose. (**A**) Protein sequences of *N. crassa* CLR-2 and *Aspergillus nidulans* ClrB homologs were used to build a phylogenetic tree using the maximum likelihood method based on the Jones-Taylor-Thornton matrix-based model using FastTree [36]. Numbers at nodes indicate the reliability of each split in the tree as computed using the Shimodaira-Hasegawa test on three alternate topologies around that split with 1,000 resamples. *R. toruloides* proteins are indicated with maroon diamonds. *N. crassa* proteins are indicated with orange squares. *A. nidulans* proteins are indicated with green circles. *S. cerevisiae* proteins are indicated with blue triangles. **(B)** Growth curves (OD600) of the indicated strains grown in yeast nitrogen base (YNB) with the indicated carbon source at 1% except for citrate which was at 8.5 mM. Lines are the average and bars or colored bands are the standard deviation of three biological replicates. *p_adj_ < 10^-4^ and **p_adj_ < 10^-10^ for *cbr1*1 vs wild type (WT) and *cbr1*1 vs *cbr1*1 + *CBR1* as determined by pairwise comparisons of estimated marginal means with a holm multiple comparison correction.

We hypothesized RTO4_13588 may play a role in regulating cellobiose utilization in *R. toruloides*. To test this hypothesis, we deleted RTO4_13588 and grew the resulting RTO4_13588Δ strain on cellobiose. Growth of cells lacking RTO4_13588 on glucose was indistinguishable from wild type cells (Fig 1B). However, deletion of RTO4_13588 resulted in a complete lack of growth on cellobiose, which could be complemented by exogenous expression of RTO4_13588 (Fig 1B). The *N. crassa* homolog of Cbr1, CLR-2, also plays a role in mannan utilization leading us to hypothesize that *CBR1* may be required for utilizing mannose, a major mannan building block [37], but *cbr1Δ* cells grew as well as wild type on mannose (Fig S2A). The closest homolog to RTO4_13588 in the model yeast *Saccharomyces cerevisiae* is the transcription factor Gal4, which regulates galactose utilization (Fig 1A) [38]. To test whether RTO4_13588 also played a role in regulating galactose utilization in *R. toruloides*, we grew wild type and RTO4_13588Δ cells with galactose as the sole carbon source. There was no detectable difference in growth between wild type cells and cells lacking RTO4_13588 on galactose (Fig S2A). Thus, we named RTO4_13588, *CBR1* for <u>c</u>ello<u>b</u>iose regulator 1.

*R. toruloides* can grow on a wide variety of soluble carbon sources [18, 31]. We asked whether *CBR1* was required for growth on any additional carbon sources. Cells lacking *CBR1* were unable to grow as well as wild type cells on the deoxy hexose sugar fucose, which could be complemented by exogenous expression of *CBR1* (Fig 1B). The growth of *cbr1*Δ cells was indistinguishable from wild type cells on all other sugars tested (fructose, xylose, arabinose, galacturonic acid, sucrose, maltose, trehalose, and raffinose) and on the lignin phenolic building blocks, coumaric and ferulic acid (Fig S2A-S2B). *CBR1* was not required for utilization of the lipid mimic Tween-80, the fermentation product acetate, or the amino acids L-alanine, proline, glutamate, or glycine, suggesting *cbr1*1 cells were capable of respiration (Fig S2C-S2D). Thus, we hypothesized *cbr1*1 cells would be able to grow on TCA cycle intermediates. To our surprise, *cbr1*1 cells had significant growth defects on citrate, α-ketoglutarate, succinate, fumarate, or malate relative to wild type and *cbr1*Δ *+ CBR1* cells (Fig 1B).

The role for Cbr1 in TCA cycle intermediate and fucose utilization could either be an expanded role for Cbr1 in *R. toruloides* relative to CLR-2/ClrB or a previously uncharacterized role for CLR-2/ClrB. To distinguish between these two possibilities, we assessed growth of the model filamentous fungus *Neurospora crassa* wild type and Δ*clr-2* cells in media containing succinate, α-ketoglutarate, malate, or fucose as the sole carbon source. Wild type *N. crassa* was only capable of limited growth on TCA cycle intermediates or fucose, and deletion of *clr-2* caused no detectable growth defect (Fig S3). These data suggest the *CBR1* role in regulating TCA cycle intermediate and fucose utilization is an expanded role for this transcription factor, relative to Cbr1 orthologs in Ascomycete filamentous fungi.

### Cbr1 activates a core regulon of five genes in response to cellobiose, TCA cycle intermediates, and fucose

We had three possible hypotheses for genetic mechanisms through which Cbr1 could regulate cellobiose, TCA cycle intermediate, and fucose utilization: (1) Cbr1 regulates genes necessary to utilize all three classes of carbon sources in response to any of these carbon sources, (2) Cbr1 regulates genes necessary to utilize each carbon source only in response to that specific carbon source, and (3) *R. toruloides* utilization of cellobiose, fucose, and/or TCA cycle intermediates proceeds through atypical metabolic pathways that require the same transporters and/or catabolic enzymes. To distinguish between these three hypotheses, we used transcriptional profiling. We exposed wild type and *cbr1*1 cells to yeast nitrogen base (YNB) lacking a carbon source (no carbon) or supplemented with 1% cellobiose, fucose, or succinate for 8 h prior to harvesting the cells for RNA sequencing (RNAseq). The cellobiose, succinate, and no carbon samples were submitted for whole transcript RNAseq, and the fucose samples were submitted for 3’ RNAseq.

Given the limited transcriptional characterization of *R. toruloides* on cellobiose or succinate, we first examined the transcriptional profile of wild type cells in these conditions relative to carbon starvation. The expression of 1168 genes was at least four-fold differentially expressed in wild type cells exposed to cellobiose compared to carbon starvation, with 155 genes upregulated and 1013 genes downregulated. In wild type cells exposed to succinate, 525 genes were at least four-fold differentially expressed relative to carbon starved cells, with 142 genes upregulated and 383 genes downregulated. Intriguingly, over 75% of genes differentially expressed on succinate (405/525) were also differentially expressed on cellobiose, supporting the hypothesis that similar sets of genes are regulated in response to cellobiose and succinate (Fig S4A-C and Dataset S1-S2).

Cells lacking *CBR1* also showed substantial transcriptional responses to cellobiose and succinate compared to carbon starvation. Five hundred seventy-nine genes were at least four-fold differentially expressed in cellobiose compared to carbon starvation. Eighty-one of the 179 genes (45%) upregulated on cellobiose in *cbr1*1 cells were also upregulated in wild type cells exposed to cellobiose, and 319 of the 400 genes (80%) downregulated on cellobiose in *cbr1*1 cells had the same expression pattern in wild type cells (Fig S4A, S4D, S4F, and Dataset S1-S2). Similarly, 742 genes were differentially expressed on succinate compared to carbon starvation in *cbr1*Δ cells. Eighty-six of the 296 genes (29%) upregulated on succinate in *cbr1Δ* cells were also upregulated in wild type cells exposed to succinate, and 256 of 446 genes (57%) downregulated on succinate in *cbr1Δ* cells were also downregulated in wild type cells exposed to succinate (Fig S4B, S4E, S4G, and Dataset S1-S2). The significant transcriptional response to cellobiose and succinate combined with the high degree of transcriptional similarity exhibited by *cbr1*1 and wild type cells suggests despite not being able to utilize cellobiose or succinate like wild type cells, *cbr1*1 cells still respond to these carbon sources to some extent.

Although many fungal transcription factor genes are upregulated in the condition in which the transcription factor plays a role, *CBR1* expression was not induced in response to cellobiose or succinate compared to carbon starvation (Fig S5A and Dataset S1-S2). To directly regulate gene expression, Cbr1 must be present in the nucleus. We hypothesized Cbr1 might be transported into the nucleus under conditions where it plays a role. To test this hypothesis, we tagged Cbr1 with green fluorescent protein (mEGFP) and the histone H2A subunit *HTA1* (RTO4_12090) with mRuby2 and exposed these cells to carbon starvation, glucose, cellobiose, or succinate to identify the localization of Cbr1. Cbr1 colocalized with the histone H2A subunit under all four conditions, indicating Cbr1 is expressed and in the nucleus during exposure to glucose, cellobiose, succinate, and carbon starvation (Fig S5B).

To gain insight into the mechanism behind the growth defects of cells lacking *CBR1*, we examined genes decreased in expression in *cbr1*Δ relative to wild type cells. Using a four-fold differential expression cutoff, 284, 18, 25, and 25 genes were downregulated in *cbr1*1 compared to wild type cells on cellobiose, succinate, carbon starvation, and fucose, respectively (Fig 2A-D and Dataset S1-S2). In concordance with our hypothesis that cellobiose, carboxylic acid, and fucose utilization are transcriptionally linked via *CBR1*, genes annotated as part of the Kyoto Encyclopedia of Genes and Genomes (KEGG) [39] 2-oxocarboxylic acid metabolism category were enriched in the *CBR1* regulon on cellobiose (Fig 2E). Carbohydrate metabolism KEGG [39] and gene ontology (GO) [40, 41] related categories were enriched on succinate, no carbon, and fucose. Further analysis revealed the carbohydrate-related enrichments primarily consisted of genes predicted to encode β-glucosidases, which cleave cellobiose into two glucose molecules (Fig 2E). Only five genes were differentially expressed by at least four-fold in at least three of the four conditions: two encoding putative β-glucosidases (RTO4_16716 and RTO4_16717) that are adjacent in the *R. toruloides* genome, two encoding putative major facilitator superfamily transporters (RTO4_10339 and RTO4_13825), and one encoding a putative phosphoglycerate mutase (RTO4_11229) (Fig 2A-D and Dataset S1-S2).

**Fig 2.**
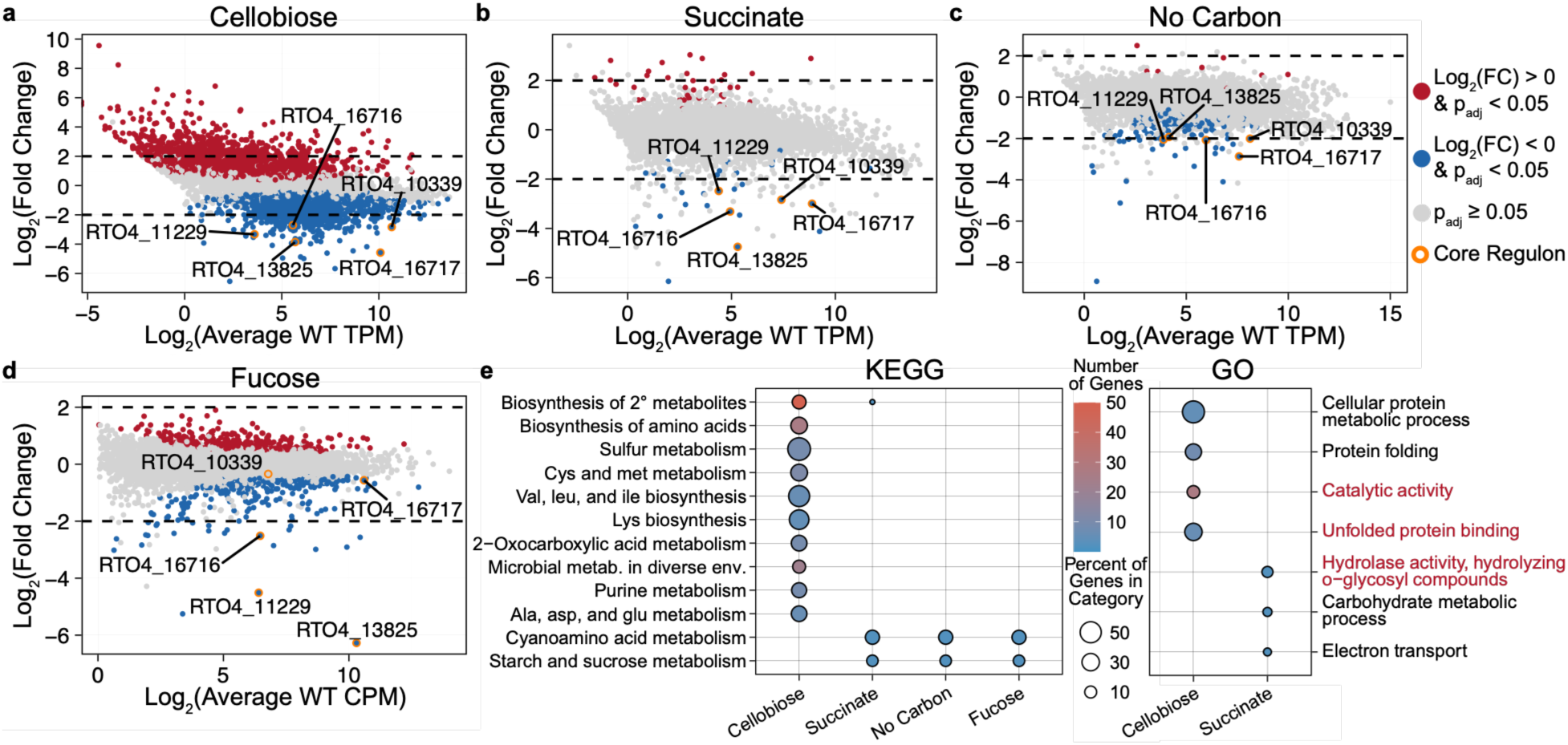
Cbr1 regulates a core regulon of five genes. (**A-D**) Scatterplots of the log_2_(average expression) of all genes in wild type (WT) cells versus the log_2_(fold change) in gene expression in *cbr1*Δ relative to wild type cells when cells were exposed to (**A**) cellobiose, (**B**) succinate, (**C**) media lacking a carbon source, or (**D**) fucose. Genes with p_adj_ < 0.05 are indicated in red or blue if they had higher or lower expression in *cbr1*Δ cells than wild type cells, respectively. Genes differentially expressed by a least four-fold in at least three of the four conditions (*CBR1* core regulon) are circled in orange and labeled with their protein IDs. Genes in grey were not significantly differentially expressed in *cbr1*Δ relative to wild type cells. Black dotted lines indicate four-fold differential expression. Standard RNAseq (**A-C**) was used to determine gene expression in transcripts per million (TPM) and fold change during exposure to cellobiose, succinate, and carbon starvation. 3’RNAseq (**D**) was used to determine gene expression in counts per million (CPM) and fold change during exposure to fucose. (**E**) Genes expressed at least four-fold lower in *cbr1*1 cells than wild type cells for each carbon source were tested for Kyoto Encyclopedia of Genes and Genomes (KEGG) [39] (left) and gene ontology (GO) [40, 41] (right) functional category enrichment. Comparisons were only displayed if they had at least one statistically significantly enriched category for each analysis. Dot color represents the number of genes differentially expressed belonging to a given category. Dot size represents the percent of genes belonging to a given category that were differentially expressed relative to all genes in that category. The text of GO biological process and molecular function categories are colored black and red, respectively. The expression of *CBR1* was left out of the scatterplots, KEGG, and GO enrichment analyses, since the change in expression of this gene was due to its deletion in the *cbr1*1 strain. All RNAseq experiments had three biological replicates.

### Cbr1-regulated β-glucosidases are necessary for cellobiose utilization

The *CBR1*-dependent expression of two predicted β-glucosidase genes led us to hypothesize that one or both β-glucosidase genes is essential for cellobiose utilization. An amino acid alignment using LALIGN [42] revealed 72% identity and 90% similarity between the two proteins, suggesting a potential for redundancy, so to test this hypothesis we deleted RTO4_16716 and RTO4_16717, together and individually. RTO4_16716Δ RTO4_167171 cells grew like wild type cells on glucose, citrate, succinate, and fucose, but were unable to grow on cellobiose (Fig 3A and Fig S6A). Analysis of the individual deletion mutants revealed that deletion of RTO4_16717 alone eliminated growth on cellobiose and complementation with RTO4_16717 restored the ability to grow on cellobiose (Fig 3B). Cells lacking RTO4_16716 alone grew as well as wild type cells on cellobiose (Fig 3B). These data indicate RTO4_16717 encodes the primary β-glucosidase for cellobiose utilization. Thus, we named RTO4_16717 and RTO4_16716, *BGL1* and *BGL2*, respectively, for <u>β</u>-<u>gl</u>ucosidase 1 and 2.

**Fig 3.**
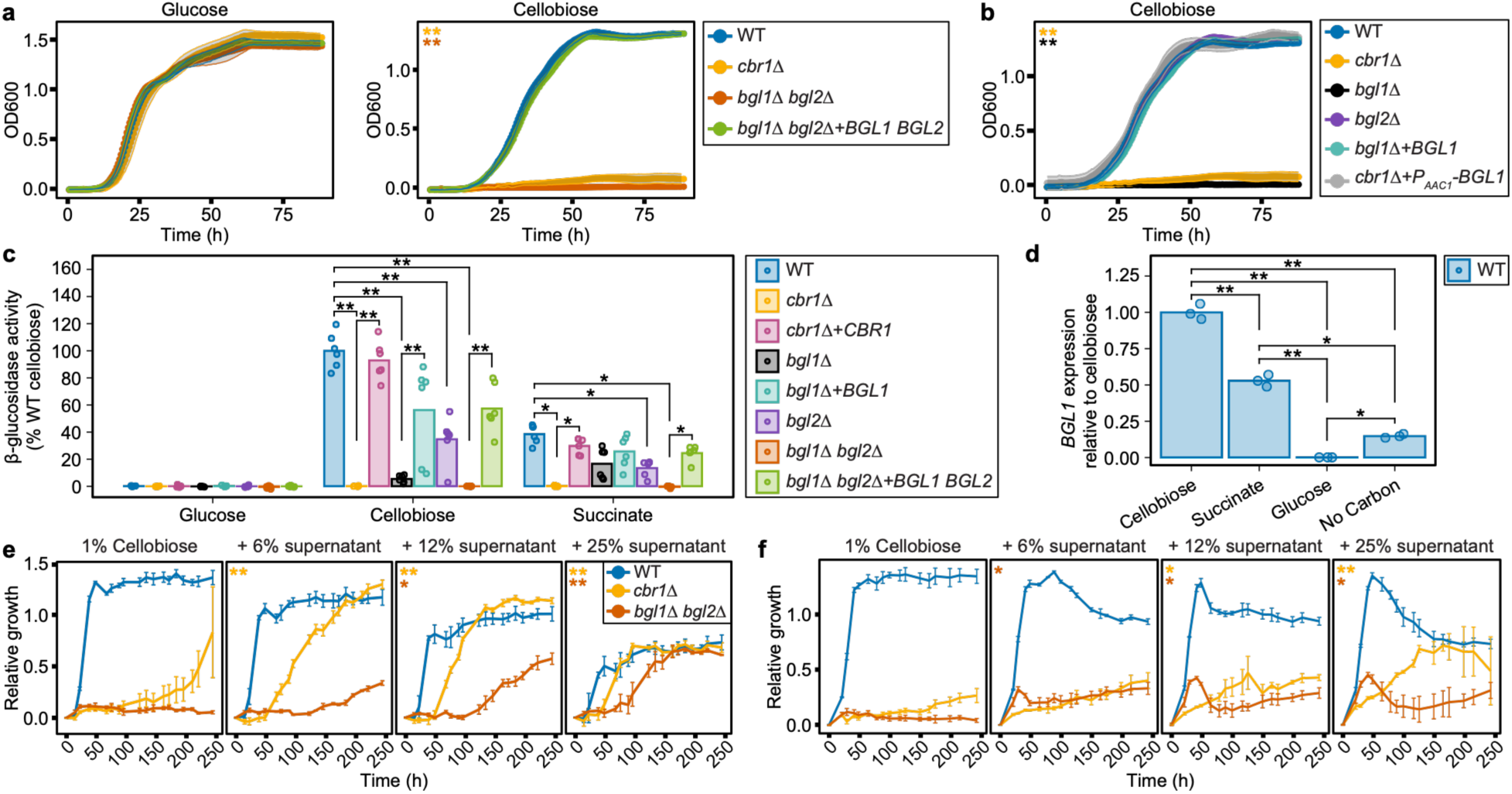
Cbr1-dependent expression of secreted β-glucosidase genes *BGL1* (RTO4_16717) and *BGL2* (RTO4_16716) is necessary for cellobiose utilization. (**A-B**) Growth (OD600) of the indicated strains in the indicated carbon sources. (**C**) β-glucosidase activity secreted by the indicated strains into culture supernatants after 20 h growth on the indicated carbon sources. β-glucosidase activity is normalized to the β-glucosidase activity of wild type (WT) cells on cellobiose. Growth (OD600) at time of supernatant harvest is in Fig S6B. (**D**) *BGL1* expression of WT cells exposed to the indicated carbon source for 4 h relative to *BGL1* expression during exposure to cellobiose for 4 h. (**E-F**) Normalized growth (OD600) of the indicated strains in 1% cellobiose supplemented with the indicated percentage of spent supernatant from wild type cells grown in (**E**) 1% cellobiose or (**F**) 1% succinate. To account for growth on nutrients remaining in the spent supernatant, growth was normalized by subtracting the growth of each replicate on media lacking a carbon source supplemented with the same concentration of spent supernatant. Growth curves without normalization, growth curves in media without a carbon source supplemented with spent supernatants, and β-glucosidase activity present in spent supernatants are in Fig S7. (**A**, **B**, **E**, and **F**) Lines are the average and bars or colored bands are the standard deviation of three biological replicates. (**C** and **D**) Bars are the average and dots are the individual data points of (**C**) five (*cbr1Δ*) or six (all other strains) biological replicates or (**D**) three biological replicates. (**A** and **B**) Colored stars indicate a significant difference in growth of the respectively colored strain compared to wild type cells and, when applicable, the respective complement strain. (**E** and **F**) Colored stars indicate a significant difference in growth of the respectively colored strain compared to growth of that strain without supernatant supplementation. *p_adj_ < 0.05 and **p_adj_ < 10^-7^ as determined by either (**A**, **B**, **E**, and **F**) pairwise comparisons of estimated marginal means with a holm multiple comparison correction, (**C**) a two-way ANOVA with a TukeyHSD post-hoc test, or (**D**) a one-way ANOVA with a TukeyHSD post-hoc test.

Both *BGL1* and *BGL2* have predicted secretion signals, leading us to hypothesize cellobiose cleavage by β-glucosidases occurs extracellularly. To address this hypothesis, we harvested supernatants of wild type, *cbr1*Δ, and *cbr1*Δ *+ CBR1* cell cultures after 20 h of growth in glucose, cellobiose, or succinate to measure secreted β-glucosidase activity. β-glucosidase activity was detected in the culture supernatant of wild type cells exposed to cellobiose indicating that β-glucosidases are indeed secreted by *R. toruloides* (Fig 3C). Consistent with our transcriptomics data, and despite the lack of a role for *BGL1* and *BGL2* in carboxylic acid utilization (Fig S6A), β-glucosidase activity was also detectable in the culture supernatant of wild type cells on succinate. Deletion of *CBR1* eliminated β-glucosidase activity in the supernatant of cellobiose and succinate cultures. This activity was restored in the supernatant of cultures of *cbr1*Δ + *CBR1* cells, indicating the secreted β-glucosidase activity was dependent on *CBR1* (Fig 3C).

Glucose cultures had no detectable β-glucosidase activity, which could either be due to reduced expression of β-glucosidase genes or reduced secretion of β-glucosidase proteins in glucose relative to cellobiose and succinate media (Fig 3C). To distinguish between these possibilities, we performed reverse transcription quantitative PCR (RT-qPCR) on *BGL1* in wild type cells. Corroborating our RNAseq data, *BGL1* expression was highest on cellobiose, followed by succinate, and then carbon starvation (Fig 3D). During exposure to glucose, *BGL1* expression was nearly undetectable, indicating the lack of β-glucosidase activity in the glucose culture supernatant was due to reduced expression of β-glucosidase genes (Fig 3C and 3D). These data may indicate β-glucosidase expression is regulated by carbon catabolite repression and that β-glucosidases potentially act as “scout” enzymes during carbon starvation similar to some polysaccharide degrading enzymes in some filamentous fungi [1, 3].

Given that *bgl1*Δ cells cannot grow when cellobiose is the sole carbon source (Fig 3B), we hypothesized Bgl1 was the source of the secreted β-glucosidase activity. To test this hypothesis, we incubated *bgl1*Δ cells in cellobiose, succinate, or glucose for 20 h. While β-glucosidase activity in the cellobiose *bgl1*Δ culture supernatant was 22-fold lower than that of wild type culture supernatants, it was detectable, and β-glucosidase activity of succinate *bgl1*Δ cultures was even more pronounced (Fig 3C and Fig S6C). This reduction in β-glucosidase secretion was mitigated in *bgl1*Δ + *BGL1* cells (Fig 3C). The low level of β-glucosidase activity remaining in *bgl1*Δ cells could either result from secretion of Bgl2 or an unidentified protein. To distinguish between these possibilities, we exposed *bgl1*Δ *bgl2*Δ and *bgl1Δ bgl2*Δ + *BGL1 BGL2* cells to cellobiose, succinate, or glucose for 20 h. Cells lacking both *BGL1* and *BGL2* had no detectable β-glucosidase activity, which was rescued by complementation with *BGL1* and *BGL2* (Fig 3C and Fig S6C). These data suggested Bgl2 contributes to β-glucosidase activity in cellobiose and succinate cultures even though *BGL2* was dispensable for growth on cellobiose. Indeed, cells lacking *BGL2* alone had a 2.8-fold reduction of β-glucosidase activity on both cellobiose and succinate relative to wild type cells, demonstrating that both Bgl1 and Bgl2 are responsible for the secreted β-glucosidase activity of *R. toruloides* (Fig 3C).

It was possible either that β-glucosidase genes were the only Cbr1-regulated genes necessary for cellobiose utilization or that other genes were also required. To distinguish between these two possibilities, we expressed *BGL1* under the constitutive *AAC1* (RTO4_12704) promoter [43] in a *cbr1*1 background (*cbr1Δ* + *P_AAC1_-BGL1*). Constitutive expression of *BGL1* in *cbr1*Δ cells resulted in wild type levels of growth on cellobiose when cells were inoculated at very low densities and a growth advantage on cellobiose relative to wild type cells at higher starting densities (Fig 3B and Fig S6D and S6E). Correspondingly, *cbr1*Δ + *P_AAC1_-BGL1* cellobiose culture supernatants had 3.2-fold higher β-glucosidase activity than wild type culture supernatants (Fig S6F).

If β-glucosidase activity is the only Cbr1-regulated requirement for cellobiose utilization, we speculated *cbr1*1 and *bgl1*1 *bgl2*1 cell growth on cellobiose could be rescued by exogenously supplied β-glucosidases. To test this hypothesis, we inoculated wild type, *cbr1*1, and *bgl1*1 *bgl2*1 cells into cellobiose supplemented with filtered, spent supernatant from wild type cells grown on cellobiose, succinate, or glucose. This spent supernatant supplementation enabled significantly improved growth of *cbr1*1 and *bgl1*1 *bgl2*1 cells, which was not due to growth on residual carbon in the spent supernatant (Fig 3E, 3F, and Fig S7). Our data demonstrate genes encoding secreted β-glucosidases, activated in response to cellobiose and TCA cycle intermediates, are the only Cbr1-regulated requirement for cellobiose utilization.

### CBR1-mediated transcriptional response to cellobiose is dependent on extracellular β-glucosidases

Extracellular degradation of cellobiose by *R. toruloides* provoked the question of whether cellobiose activates the *CBR1*-dependent transcriptional response to cellobiose or if the activating signal is a cellobiose catabolic derivative. We differentiated between these possibilities in two ways.

We first compared the transcriptional profile of *bgl1*1 *bgl2*1 cells exposed to cellobiose, which have no extracellular β-glucosidase activity but contain wild type copies of *CBR1*, to that of wild type and *cbr1*Δ cells using 3’ RNAseq. The transcriptional profiles of *bgl1*1 *bgl2*1 cells were more similar to *cbr1*1 than wild type cells (Fig 4A-C and Fig S8 and Dataset S3-S4).

**Fig 4.**
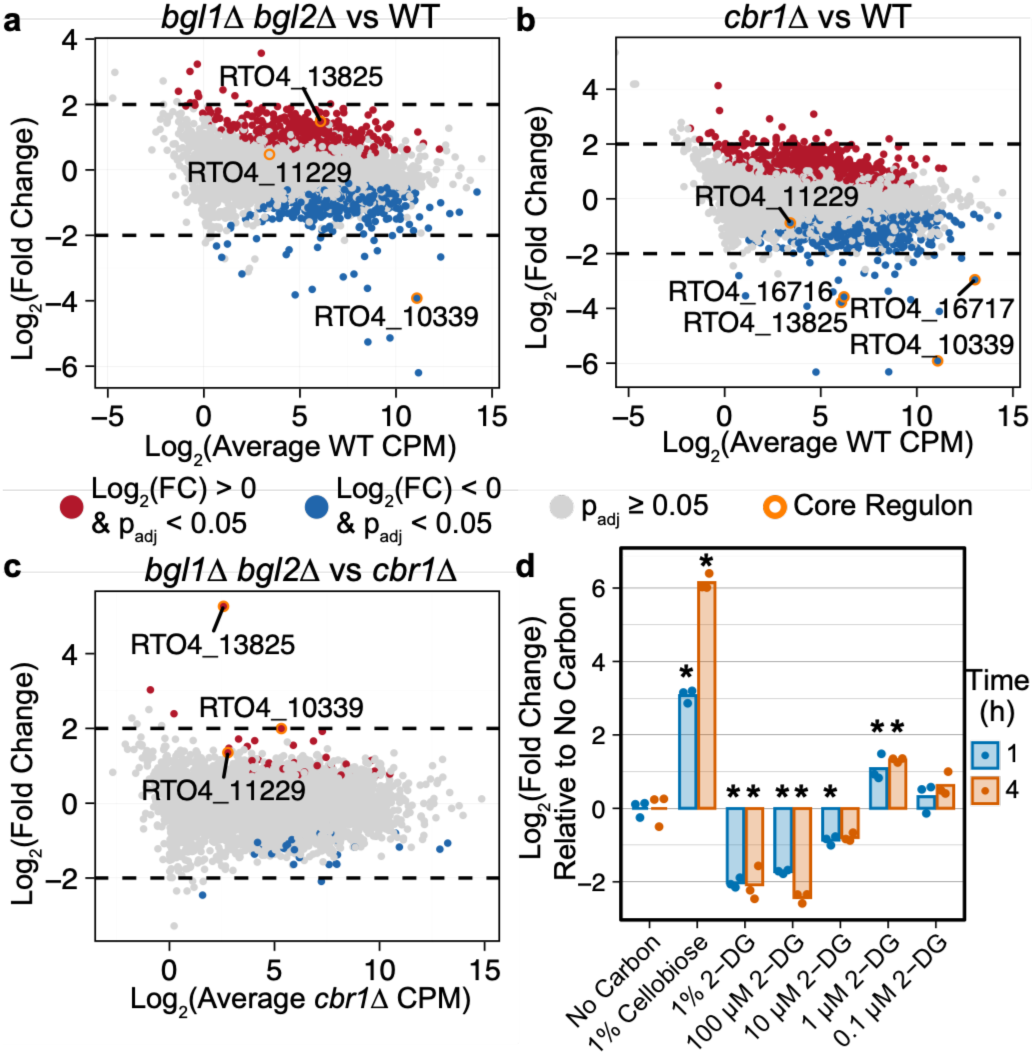
*CBR1*-mediated transcriptional response to cellobiose is dependent on secreted β-glucosidases. (**A-C**) Scatterplots of the log_2_(average expression) of all genes in (**A-B**) wild type (WT) or (**C**) *cbr1*Δ cells versus the log_2_(fold change) in gene expression for the indicated comparison when cells were exposed to cellobiose as measured by 3’RNAseq. Genes with p_adj_ < 0.05 are indicated in red or blue if they had higher or lower expression, respectively, in (**A** and **C**) *bgl1*1 *bgl2*1 cells or (**B**) *cbr1*Δ cells. The genes in the *CBR1* core regulon are circled in orange and labeled with their respective protein IDs. The expression of *CBR1*, *BGL1,* and *BGL2* were left out of the scatterplots involving their respective deletion strains. RNAseq experiments had three biological replicates. (**D**) *BGL1* expression in wild type cells exposed to carbon starvation for 1 h followed by exposure to continued carbon starvation or the indicated concentrations of cellobiose or 2-deoxyglucose (2-DG) as the sole carbon source for 1 h or 4 h. Expression values were normalized to the average expression of *BGL1* on no carbon for each timepoint and log_2_ transformed. Bars are the average and dots are the values of three biological replicates. *p_adj_ < 0.05 relative to the no carbon condition for the respective time point, as determined by two-way ANOVA with a TukeyHSD post-hoc test.

Thirty-five genes were at least four-fold differentially expressed in *bgl1*1 *bgl2*1 relative to wild type cells (12 upregulated and 23 downregulated) (Fig 4A and Dataset S3-S4). Twenty-three of these genes were also at least four-fold differentially expressed in *cbr1*1 compared to wild type cells (Fig 4B and Dataset S3-S4). In contrast, only five genes were at least four-fold differentially expressed in *bgl1*1 *bgl2*1 relative to *cbr1*1 cells, which included a 38-fold upregulation of the *CBR1*-regulated predicted transporter gene RTO4_13825 (Fig 4C and Dataset S3-S4). The expression pattern of RTO4_13825, raised the question of whether any other genes in the core Cbr1 regulon were involved in cellobiose sensing or utilization. Cells lacking RTO4_13825, RTO4_10339, or RTO4_11229 all grew as well as wild type on cellobiose (Fig S9). These data indicate the signal for the *CBR1*-dependent cellobiose response requires secreted β-glucosidases, suggesting cellobiose is likely not the signaling molecule for Cbr1 activation.

β-glucosidases cleave cellobiose into two glucose molecules. Thus, a second way we answered the question of whether cellobiose or a cellobiose catabolic derivative is the activation signal was to determine whether low glucose concentrations activated Cbr1-regulated gene expression. To mitigate changes in glucose concentration due to metabolism, we used the non-metabolizable glucose analog 2-deoxyglucose. We exposed wild type cells to 10-fold dilutions of 2-deoxyglucose (100 μM to 0.1 μM), 1% 2-deoxyglucose, 1% cellobiose, or carbon starvation for 1 h and 4 h and measured *BGL1* transcription via RT-qPCR. As expected, cells exposed to cellobiose expressed *BGL1* at high levels, and *BGL1* expression was repressed during exposure to 1% 2-deoxyglucose (Fig 4D). Exposure to 1 μM 2-deoxyglucose resulted in a two to three-fold increase in *BGL1* expression relative to carbon starvation at both timepoints (Fig 4D). This *BGL1* activation did not rise to the same magnitude as exposure to cellobiose itself, perhaps due to differences in glucose concentration, the rate glucose enters the cell, or a lack of resources due to starvation on 2-deoxyglucose. However, increased *BGL1* expression on 1 μM 2-deoxyglucose suggests a low concentration of glucose, a catabolic cellobiose derivative, activates β-glucosidase expression.

### A Cbr1-regulated transporter gene is necessary for citrate utilization

Two genes encoding predicted major facilitator superfamily transporters (RTO4_13825 and RTO4_10339) and a gene encoding a predicted phosphoglycerate mutase (RTO4_11229) were also members of the *CBR1* core regulon. We hypothesized these genes may be necessary for growth on TCA cycle intermediates and/or fucose. To test this hypothesis, we grew RTO4_13825Δ, RTO4_103391, and RTO4_11229Δ cells on media containing glucose, fucose, citrate, succinate, α-ketoglutarate, malate, or fumarate. The growth of RTO4_10339Δ and RTO4_11229Δ cells was indistinguishable from wild type cells under all conditions tested (Fig S9). Cells lacking RTO4_13825 were unable to grow on citrate but grew as well as wild type cells on all other carbon sources tested (Fig 5A and Fig S9A). Thus, we named RTO4_13825 *TCT1* for tri<u>c</u>arboxylic acid transporter.

**Fig 5.**
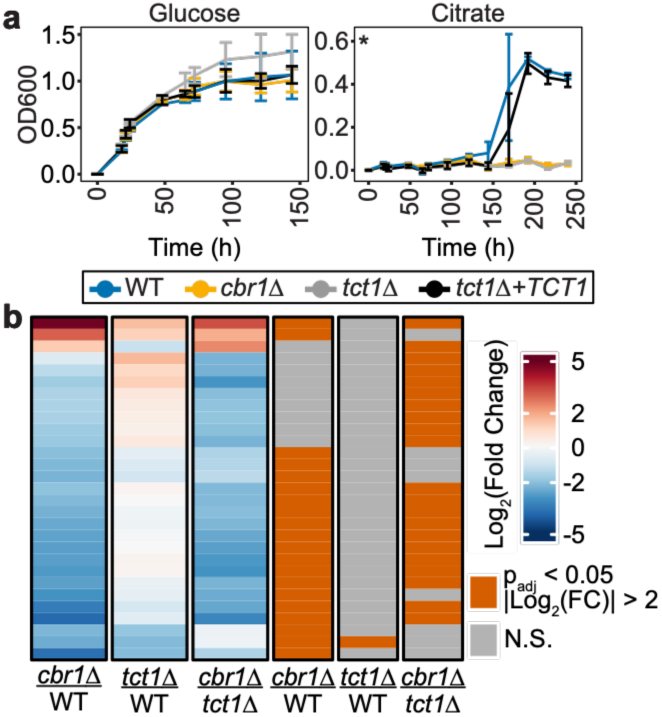
*TCT1* (RTO4_13825) is required for citrate utilization. (**A**) Growth (OD600) of the indicated strains in YNB 1% glucose or 8.5 mM citrate. Lines are the average and bars are the standard deviation of three biological replicates. *p_adj_ < 10^-7^ of *cbr1*Δ vs wild type (WT), *tct1*Δ vs WT, and *tct1*Δ vs *tct1Δ + TCT1* as determined by pairwise comparisons of estimated marginal means with a holm multiple comparison correction. (**B**) Heatmap of the log_2_(fold change) in the expression of genes at least four-fold differentially expressed in one or more of the indicated comparisons when cells were exposed to citrate as the sole carbon source, as measured by 3’RNAseq. Orange bars to the right of the heatmap indicate significance using the criteria of p_adj_ < 0.05 and at least four-fold change in expression, while grey bars indicate genes that do not meet those criteria in the indicated comparison (N.S., not significant). The expression of *CBR1* and *TCT1* were left out of the heatmaps since changes in expression were due to their respective deletions in the *cbr1*1 and *tct1*1 strains. RNAseq experiments had three biological replicates.

To gain further insight into the role of *TCT1*, we conducted RNAseq on wild type, *cbr1*1, and *tct1*1 cells exposed to citrate for 24 h. Only 20 genes were at least four-fold differentially expressed in *cbr1*Δ relative to wild type cells (Fig 5B and Dataset S5-S6). Surprisingly, these 20 genes did not include *TCT1* or genes predicted to play a role in citrate metabolism. Of the other genes in the *CBR1* core regulon, *BGL1* and *BGL2* were downregulated by 6.7 and 3.2-fold, respectively, in *cbr1*Δ relative to wild type cells, but RTO4_10339 and RTO4_11229 were not differentially expressed (Fig 5B and Dataset S5-S6). Only one gene, RTO4_15140, had decreased expression in *tct1*Δ cells relative to wild type cells. RTO4_15140 was also downregulated in *cbr1*1 relative to wild type cells (Fig 5B and Dataset S5-S6). RTO4_15140 is predicted to encode a P-type ATPase transduction domain A containing protein with a single transmembrane domain, and any potential role in citrate utilization is unclear. The similarity of *tct1*1 and wild type cell transcriptional profiles may suggest *tct1*1 cells can sense citrate, despite not being able to grow.

### Cbr1 inhibits carbon catabolite repression during glucose-glucose disaccharide utilization

Glucose is a highly preferred carbon source in most fungi. Extracellular β-glucosidases cleave cellobiose into two glucose molecules, which we hypothesized would repress cellobiose utilization genes via carbon catabolite repression [1, 3, 4]. *BGL1* expression is almost undetectable when wild type cells are exposed to 1% glucose or at least 100 µM 2-deoxyglucose (Fig 3D and 4D). Thus, we hypothesized wild type cell growth on cellobiose would be inhibited by the non-hydrolyzable glucose analog 2-deoxyglucose. Indeed, 61 µM 2-deoxyglucose fully inhibited *R. toruloides* growth on 1% cellobiose, indicating glucose is preferred over cellobiose by *R. toruloides* (Fig 6A).

**Fig 6.**
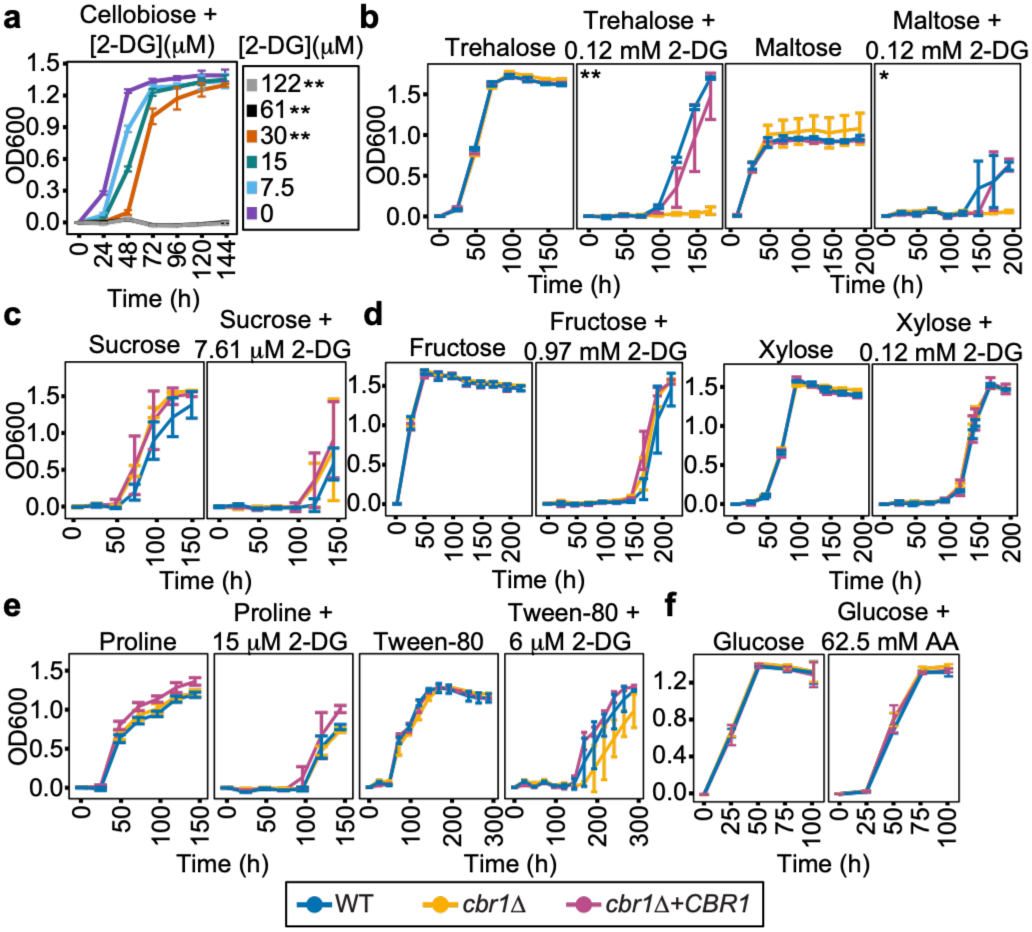
Cbr1 inhibits carbon catabolite repression during glucose-glucose disaccharide utilization. (**A**) Growth (OD600) of wild type (WT) cells in 1% cellobiose supplemented with the indicated concentration of 2-deoxyglucose (2-DG). (**B-F**) Growth of the indicated strains on the indicated carbon sources with the indicated concentration of 2-DG or allyl alcohol (AA). Lines are the average and bars are the standard deviation of (**A-E**) three or (**F**) four biological replicates. *p_adj_ < 0.01 and **p_adj_ < 10^-4^ for (**A**) 1% cellobiose supplemented with the indicated 2-DG concentration vs 1% cellobiose or (**B-F**) *cbr1*Δ vs WT and *cbr1*Δ vs *cbr1*Δ + *CBR1* as determined by pairwise comparisons of estimated marginal means with a holm multiple comparison correction.

This carbon preference represents a biological challenge during utilization of substrates in which glucose is released during catabolism; a negative feedback loop could repress substrate utilization genes in response to the released glucose, slowing growth on the carbon source. We hypothesized *CBR1* may inhibit carbon catabolite repression to bypass this negative feedback loop. To test this hypothesis, we grew wild type, *cbr1*1, and *cbr1*1 + *CBR1* cells on a nonpreferred carbon source and 2-deoxyglucose. Cells lacking *CBR1* were significantly more sensitive to 2-deoxyglucose than wild type or *cbr1*Δ + *CBR1* cells when the glucose-glucose disaccharides maltose or trehalose were the carbon source, suggesting Cbr1 is a negative regulator of carbon catabolite repression (Fig 6B).

Canonically, regulators of carbon catabolite repression regulate a broad spectrum of nonpreferred carbon source utilization genes [1, 3–10]. We hypothesized *CBR1* would play a similar role in *R. toruloides*. However, the growth of *cbr1*Δ cells exposed to 2-deoxyglucose in combination with the glucose-fructose disaccharide sucrose, the hexose monosaccharide fructose, the pentose monosaccharide xylose, the amino acid proline, or the lipid mimic Tween-80 was indistinguishable from wild type cells (Fig 6C-6E). To measure the role of *CBR1* in regulating genes involved in alcohol utilization, we used allyl alcohol, which is converted to the toxic molecule acrolein by alcohol dehydrogenases. Alcohol dehydrogenase expression is repressed by glucose [44]. Wild type and *cbr1*1 cells were equally sensitive to allyl alcohol, indicating glucose-mediated repression of alcohol dehydrogenases is not dependent on Cbr1 (Fig 6F). To determine whether any *CBR1* core regulon genes play a role in carbon catabolite repression, we tested 2-deoxyglucose sensitivity of *bgl1*1 *bgl2*1, *tct1*1, RTO4_10339Δ, and RTO4_112291 cells. The sensitivity of these strains to 2-deoxyglucose during exposure to trehalose, maltose, sucrose, proline, xylose, fructose, and tween-80 was indistinguishable from wild type cells (Fig S10). These data indicate Cbr1 inhibits carbon catabolite repression of glucose-glucose disaccharides either by directly regulating gene expression or through genes outside of the *CBR1* core regulon.

A role for Cbr1 Ascomycete homologs in carbon catabolite repression regulation has not been identified, so we asked whether the *N. crassa* Cbr1 homolog CLR-2 plays a similar role. The sensitivity of wild type and Δ*clr-2* cells to 2-deoxyglucose was indistinguishable during growth on the glucose-glucose disaccharide trehalose or the monosaccharide xylose (Fig S11). These data indicate the role for Cbr1 in *R. toruloides* in inhibiting carbon catabolite repression of glucose-glucose disaccharides is an expanded role for Cbr1 relative to the model Ascomycete ortholog CLR-2.

## DISCUSSION

Gene regulation in response to nutrient availability requires cells to integrate information from activating and repressing nutrient signals. Previously described fungal transcriptional networks regulating carbon utilization are made up of carbon source-specific transcription factors that activate genes necessary to utilize a particular carbon source when it is present and broad-acting transcription factors that regulate carbon catabolite repression [1, 2]. However, studies of these carbon source utilization transcriptional networks have mainly focused on Ascomycete fungi. We investigated nutrient sensing regulatory mechanisms in a Basidiomycete yeast and identified a transcription factor that coregulates carbon source-specific gene activation and carbon catabolite repression specifically of glucose-glucose disaccharide utilization genes (Fig 7).

**Fig 7.**
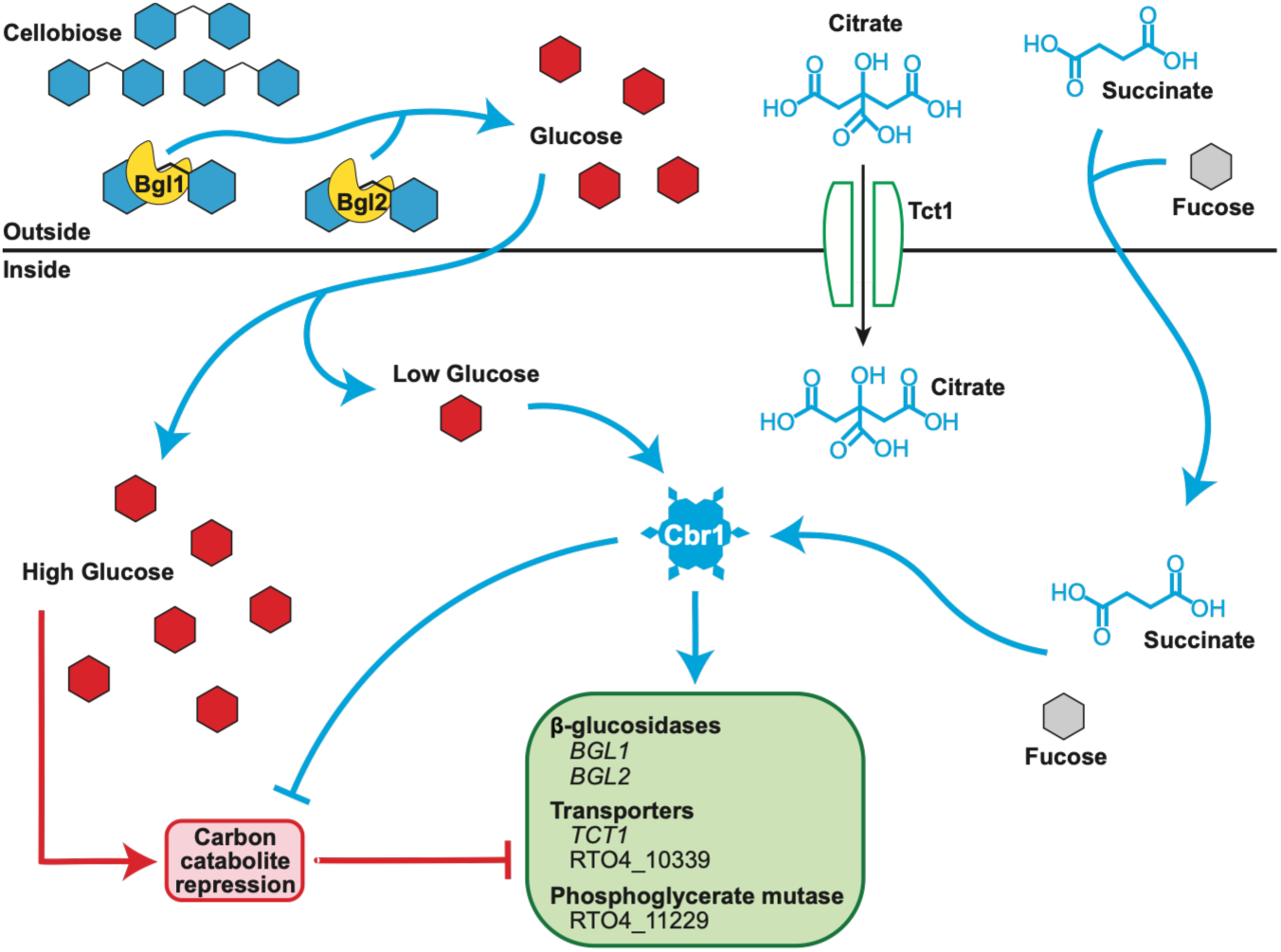
Model for Cbr1-mediated regulation of cellobiose, TCA cycle intermediate, and fucose utilization and carbon catabolite repression. Cbr1 is activated by the presence of a low glucose concentration, fucose, and TCA cycle intermediates or their metabolic derivatives. Activated Cbr1 induces the expression of genes encoding β-glucosidases, predicted transporters, and a predicted phosphoglycerate mutase. The Cbr1-regulated predicted transporter gene *TCT1* is required for citrate utilization. Tct1 is placed at the plasma membrane in this model, but the localization of Tct1 is currently unknown and should be the subject of future studies. Transporters for other TCA cycle intermediates and fucose are unknown. The β-glucosidase genes *BGL1* and *BGL2* encode secreted β-glucosidases that cleave cellobiose into two glucose molecules. *BGL1* is required for growth on cellobiose. A high glucose concentration activates carbon catabolite repression, which represses β-glucosidase gene expression and likely represses the expression of genes necessary to utilize fucose and TCA cycle intermediates. Cbr1 plays a role in inhibiting carbon catabolite repression, specifically when a glucose-glucose disaccharide is present as the nonpreferred carbon source.

### Cbr1 regulation of cellobiose, TCA cycle intermediate, and fucose utilization may shed light on complex microbial interactions

There have been limited investigations of genetic regulators of cellulose and cellobiose utilization in Basidiomycetes. A recent study identified Roc1 as a regulator of cellulase-encoding genes in the basidiomycete wood-decaying fungus *Schizophyllum commune* [45]. However, homologs of Roc1 do not exist outside of the Agaricomycetes [45]. In *Ganoderma lucidum*, the R2R3-type MYB transcription factor *Gl*Myb activates cellulase gene expression [46]. In Ascomycete filamentous fungi, CLR-2/ClrB regulates cellulose degradation and utilization through direct activation of cellulase genes [32, 47]. We identified the CLR-2/ClrB ortholog Cbr1 in the oleaginous, basidiomycete yeast *R. toruloides* (Fig 1). Although Ascomycete and Basidiomycete fungi are separated by approximately 400 million years of evolution [12], Cbr1 is required to utilize the cellulose breakdown product cellobiose (Fig 1). *N. crassa* CLR-2 does not directly regulate the expression of genes encoding β-glucosidases, which are regulated by CLR-1 [32, 47]. However, although the role of Cbr1 is significantly expanded beyond that of CLR-2/ClrB to include inhibition of carbon catabolite repression and utilization of TCA cycle intermediates and fucose, Cbr1 regulation of cellobiose utilization indicates some functional conservation of CLR-2/ClrB/Cbr1 exists between Ascomycete filamentous fungi and Basidiomycete yeast.

Cbr1 regulates citrate utilization through the predicted transporter gene *TCT1* and cellobiose utilization through the β-glucosidase genes *BGL1* and *BGL2* (Fig 7). Genetic mechanisms through which Cbr1 regulates utilization of fucose and dicarboxylic acid TCA cycle intermediates remains unclear. It is possible the two uncharacterized genes in the *CBR1* core regulon, the predicted transporter RTO4_10339 and the predicted phosphoglycerate mutase RTO4_11229, play roles in fucose and/or dicarboxylic acid utilization that are redundant with other *CBR1*-regulated genes that did not meet our strict *CBR1* core regulon criteria (Fig 2 and Fig S4). It will be the role of future studies to characterize roles for RTO4_10339, RTO4_11229, and other genes that may be responsible for fucose and dicarboxylic acid catabolism in the *CBR1* pathway.

In other fungal species, utilization of multiple carbon sources is occasionally regulated by a single transcription factor. CLR-2 in *N. crassa* regulates cellulose utilization, mannan utilization, and a small number of hemicellulose utilization genes [3, 32, 37, 47]. In *Trichoderma reesei*, *Penicillium oxalicum*, and some Aspergilli, the transcription factor Xyr1/XlnR regulates cellulose and hemicellulose utilization [33–35]. ARA-1/Ara1 regulates arabinan and galactose utilization in *N. crassa* and *T. reesei* [3, 48]. FarA regulates utilization of fatty acids and the lignin component ferulic acid in Aspergilli [49, 50]. In *Candida albicans*, Adr1 is required for growth on citrate and substrates that feed into the TCA cycle, like glutamate and malate [51]. However, in these examples linkage of these carbon sources can be explained either by the presence of both carbon sources in the plant cell wall (i.e., cellulose and hemicellulose or arabinan and galactose) or shared catabolic pathways (i.e., fatty acids and ferulic acid or citrate, glutamate, and malate).

Cellobiose, fucose, and TCA cycle intermediates are not linked by known catabolic pathways, and our data do not support catabolism through the same noncanonical metabolic pathway in *R. toruloides*. Cbr1-regulated β-glucosidase genes are critical for cellobiose but not fucose or TCA cycle intermediate utilization (Fig 3 and Fig S6A), while Tct1 is necessary for citrate but not cellobiose or fucose utilization (Fig 5A and Fig S9A). Aside from *CBR1*, we did not identify genes necessary for utilization of all three of cellobiose, fucose, and TCA cycle intermediates.

We speculate coregulation of cellobiose, fucose, and TCA cycle intermediates may suggest *R. toruloides* encounters cellobiose, fucose, and TCA cycle intermediates together in nature. One possible explanation is *R. toruloides* scavenges cellobiose and fucose during plant biomass degradation by filamentous fungi, which secrete organic acids like citrate or succinate to acidify their environment and can contain fucose in their cell walls [52–54]. β-glucosidase gene activation that is primed by organic acids secreted by fungi or fucose in fungal cell walls before encountering cellobiose released from cellulose could confer a growth advantage to *R. toruloides* over microbes that only activate β-glucosidase genes in response to cellobiose itself. Indeed, high densities of cells constitutively expressing *BGL1* that do not require time to activate β-glucosidase genes had a growth advantage over wild type cells that activated β-glucosidase genes in response to cellobiose (Fig S6D).

### Cbr1 regulation of carbon catabolite repression specifically of glucose-glucose disaccharides disrupts a negative feedback loop for disaccharide utilization

Carbon catabolite repression enables prioritization of carbon sources that require less energy to metabolize over carbon sources that require more energy to metabolize [1]. Known carbon catabolite repression genetic mechanisms regulate genes required for utilization of all less preferred carbon sources when a more preferred carbon source is present [3–10]. In contrast, Cbr1 inhibits carbon catabolite repression specifically of glucose-glucose disaccharide utilization (Fig 6). We speculate this mechanism allows continued expression of genes that hydrolyze disaccharide bonds even when this hydrolysis produces preferred monosaccharides, combating a negative feedback loop (Fig 7). To our knowledge, genetic linkage via a single transcription factor of carbon catabolite repression inhibition with activation of genes necessary for nonpreferred carbon source utilization has not been described in fungi. Additionally, regulation of genes involved in utilization of a specific class of nonpreferred carbon sources in response to a preferred carbon source represents a previously uncharacterized form of carbon catabolite repression regulation.

Fungal pathogens of plants and animals that do not regulate carbon catabolite repression like wild type cells frequently have reduced virulence [8, 55–57]. Carbon utilization and carbon catabolite repression regulation by fungi also affects how they respond to antifungal drugs [14, 15]. Thus, understanding diverse carbon catabolite repression mechanisms present throughout the fungal kingdom is critical in controlling fungal disease. Additionally, a more complete understanding of mechanisms of carbon catabolite repression and the diversity of nutrient sensing transcriptional networks is necessary for harnessing the primary and secondary metabolic capabilities of the fungal kingdom for biotechnology. Mapping these networks will be critical in metabolically engineering fungi for green industrial fermentation.

## MATERIALS and METHODS

### Strains and culturing

*R. toruloides* strains used in this study are listed in Table S1. All strains were derived from the wild type reference strain IFO 0880 (also called NBRC 0880, obtained from Biological Resource Center, NITE (NBRC), Japan). The starting strain for genetic manipulations was the non-homologous end-joining deficient *yku70*1 (RTO4_11920Δ) strain [58]. *R. toruloides* was transformed using *Agrobacterium tumefaciens*-mediated transformation as previously described [26], using *A. tumefaciens* EHA 105 and plasmids derived from pGI2 [59]. All strains were confirmed by PCR and/or DNA sequencing of transformed loci. For details see Text S1. *R. toruloides* strains were grown from freezer stocks on yeast extract peptone dextrose (YPD) (VWR) + 2% agar (BD Difco™ Bacto™) plates at 28°C for 2-3 d prior to inoculation into liquid YPD to grow strains at 28°C with constant shaking at 250 rpm to exponential phase growth prior to starting experiments.

*N. crassa* strains used in this study are listed in Table S2. These strains were derived from the wild type reference strain FGSC 2489 and obtained from the Fungal Genetics Stock Center [60, 61]. *N. crassa* cells were grown from freezer stocks on Vogel’s minimal medium [62] + 2% sucrose + 1.5% agar (BD Difco™ Bacto™) slants for 2 d at 28°C in the dark and 4 d at 28°C in constant light. *N. crassa* conidia were harvested from slants with sterile double distilled H_2_O for inoculation.

All *R. toruloides* experiments were performed with yeast nitrogen base (YNB) with NH_4_SO_4_ without amino acids (MP Biomedicals) + 12 mM K_2_HPO_4_ (Fisher) + 38 mM KH_2_PO_4_ (Fisher) + 100 nM Fe(II)SO_4_ pH 6.2 with the indicated carbon source at 1% w/v with the exception of coumaric acid and ferulic acid, which were added at a concentration of 0.2% w/v, and citrate which was added at a concentration of 8.5 mM. All *N. crassa* experiments were performed using Vogel’s minimal medium [62] with 50 mM NH_4_Cl instead of 25 mM NH_4_NO_3_ (VMM NH_4_Cl) and the indicated carbon source at 1% w/v. Carbon sources used are listed in Table S3.

### R. toruloides and N. crassa growth experiments

*R. toruloides* colonies were inoculated into liquid YPD and grown at 28°C with constant shaking at 250 rpm to exponential phase. Cells were then washed and inoculated into 750 μL of the indicated media in 48-well plates at 0.01 OD600/mL or 0.1 OD600/mL, except for the growth curve in Fig S6D, which was inoculated at 0.5 OD600/mL as measured by an Eppendorf BioPhotometer with a 10 mm path length. Plates were incubated at 28°C with constant shaking (3 mm orbit, 600 rpm for individual time points and 3 mm orbit, 425 rpm for kinetic growth experiments). Growth was measured as OD600 on an Agilent BioTek Synergy HTX for individual time points or an Agilent BioTek Epoch2 for kinetic growth experiments. For details see Text S1.

*N. crassa* conidia were inoculated into either 100 mL of liquid VMM NH_4_Cl with the indicated carbon source in 250 mL flasks or 3 mL of liquid VMM NH_4_Cl with the indicated carbon source in round-bottomed, deep-well 24-well plates. After 24 h (flasks) or 7-10 d (24-well plates) biomass was harvested, dried for 24 h in a 60°C drying oven, and weighed. For details see Text S1.

### Phylogenetic tree generation

The Basic Local Alignment Search Tool for proteins (BLASTP) was used to identify homologs of CLR-1, CLR-2, and XLR-1 from *N. crassa* (OR74A) and ClrA, ClrB, and XlnR from *Aspergillus nidulans* (A4) in the proteomes of *N. crassa*, *A. nidulans*, *S. cerevisiae* (S288C), and *R. toruloides* (IFO 0880). CLR-1, CLR-2, and XLR-1 from *N. crassa* (OR74A) and ClrA, ClrB, and XlnR from *A. nidulans* (A4) protein sequences were obtained from Mycocosm [63]. Homologs with E < 10^-10^ were aligned via MUSCLE [64] in MEGA11 [65] and the phylogenetic tree was generated using FastTree [36] using the maximum likelihood method based on the Jones-Taylor-Thornton matrix-based model and visualized in MEGA11. The reliability of each split in the tree was computed using the Shimodaira-Hasegawa test on three alternate topologies around that split. Trees were visualized in MEGA11 [65] and rooted on the midpoint.

### Microscopy

Fluorescent microscopy was performed using a Nikon Eclipse Ti2-E inverted microscope equipped with a prime BSI express CMOS camera using a CFI Plan Apochromat 100X oil objective. For details see Text S1.

### RNA sequencing and transcript abundance

Flasks containing 100 mL of YNB without a carbon source or YNB with the indicated carbon source were inoculated with the indicated exponentially growing cells at 0.1 OD600/mL (measured by an Eppendorf BioPhotometer). Culture conditions and sequencing methods for each dataset are summarized in Table S4. Cells were harvested and flash frozen in liquid N_2_. RNA was extracted using the Quick-RNA Fungal/Bacterial Miniprep Kit (Zymo), followed by either treatment with the TURBO DNA-free kit (Invitrogen) for standard RNAseq or DNase I treatment (NEB) and purification with the Monarch^®^ Spin RNA Cleanup Kit (NEB) for 3’RNAseq. Standard RNAseq was performed at the California Institute for Quantitative Biosciences at the University of California Berkeley (QB3-Berkeley) on an Illumina NextSeq 2000 with 150 bp paired end reads. 3’RNAseq was performed at the Biotechnology Resource Center at Cornell University on an Illumina NovaSeqX with 150 bp paired end reads.

Reads were aligned to the *R. toruloides* IFO 0880 genome (v4) [26]. For standard RNAseq, transcript abundance (transcripts per million, TPM) was quantified using Salmon v. 1.10.0 [66]. For 3’RNAseq, reads were aligned using STAR v. 2.7.11b [67], counted with HTSeq v. 2.0.9 [68], and counts per million (CPM) values were calculated by dividing counts for a gene by total counts for the sample and multiplying by 10^6^. The GFF file used to analyze 3’RNAseq data is available in Dataset S7. Differential expression was determined using DESeq2 v. 1.50.1 [69]. RNAseq data were deposited in the Gene Expression Omnibus (GEO) at the National Center for Biotechnology Information (NCBI) and are accessible through GEO series accession numbers GSE293943 (standard RNAseq) and GSE313844 (3’RNAseq). For details see Text S1.

### Functional enrichment analysis

KEGG and GO enrichment analyses were performed using the enricher function of the clusterProfiler R package v. 4.18.1 with Benjamini-Hochberg multiple hypothesis testing correction [70]. For details see Text S1.

### Reverse transcription quantitative PCR

To measure *BGL1* expression, wild type cells were exposed to the indicated carbon source in 100 mL of the indicated media in 250 mL flasks and incubated at 28°C with shaking at 200 rpm for the indicated length of time. Cells were harvested, flash frozen in liquid N_2_, and RNA was purified as described above. RT-qPCR was performed using the Luna^®^ Universal One-Step RT-qPCR Kit (NEB) on a CFX96^TM^ Real-Time System (Bio-Rad). For details see Text S1.

### β-glucosidase enzyme activity assays

β-glucosidase activity was measured using p-nitrophenyl-β-D-glucopyranoside (EMD Millipore Corp.). For details see Text S1.

### Spent media growth assays

Filtered supernatants of wild type cells growing in YNB 1% glucose, YNB 1% cellobiose, or YNB 1% succinate were added to YNB lacking a carbon source or YNB 1% cellobiose at the indicated concentration. The indicated strains were inoculated into this media at 0.01 OD600/mL (measured by an Eppendorf BioPhotometer) in 750 μL in 48-well plates and incubated at 28°C with shaking at 600 rpm (3 mm orbit). For details see Text S1.

### Statistical significance tests

All statistical analyses were performed in Rstudio (v2025.09.2+418; R v4.5.2) using either a one-way or two-way ANOVA with a Tukey post-hoc test (bar graphs), a two-tailed Welch’s two-sample *t*-test (bar graphs), or a general linear model with two-tailed post-hoc analysis using the emmeans function of the emmeans R package [71] with a holm multiple comparison correction (line graphs). Statistical significance for differential expression of RNAseq data was determined using DESeq2 [69]. All experiments were performed with a minimum of three biological replicates. The exact number of replicates for each experiment is noted in the relevant figure legend.

## Data availability

Raw RNAseq data is available in GEO at the NCBI through accession numbers GSE293943 (standard RNAseq) and GSE313844 (3’RNAseq). Processed RNAseq data is available in Dataset S1-S6. The numerical values used to generate all other graphs are in Dataset S9. Strains constructed in this study are available upon request.

## ACKNOWLEDGEMENTS

We thank members of the Huberman lab for helpful comments on the manuscript. This work was supported by a grant from the National Institute of General Medical Sciences of the National Institutes of Health (NIH) under Award Number R35GM150926 to L.B.H. J.D.K. was supported by NIH Ruth L. Kirschstein National Research Service Award Institutional Research Training Grant 5T32AI145821-03. This work utilized the Cornell Institute of Biotechnology Biotechnology Resource Center Genomics Facility (RRID:SCR_021727) and BioHPC (RRID:SCR_021757) and the QB3 Genomics core at the University of California Berkeley (RRID:SCR_022170).

