## Supplementary material for "Carbohydrate utilization regulator reveals a noncanonical mechanism of nutrient differentiation": SI Appendix

**SUPPORTING INFORMATION**

**Text S1. SI MATERIALS and METHODS**

*Rhodotorula (Rhodosporidium) toruloides strain construction*

*R. toruloides* strains used in this study are listed in Table S1. All strains were derived from the wild type reference strain IFO 0880 (also called NBRC 0880, obtained from Biological Resource Center, NITE (NBRC), Japan). The starting strain for genetic manipulations was the non-homologous end-joining deficient *yku70*Δ (RTO4_11920Δ) strain [1]. *R. toruloides* was transformed using *Agrobacterium tumefaciens*-mediated transformation as previously described [2], using *A. tumefaciens* EHA 105 and plasmids derived from pGI2 [3]. Most gene deletions, complementations, constitutive expressions, and tags were performed using homologous recombination of a nourseothricin or G418 resistance cassette with flanking arms of ~1,000 bp. Nourseothricin (GoldBio) and G418 (VWR) were both added to selection media at 100 μg/mL.

There were a few specialized cases of strain construction. We fluorescently tagged the histone H2A subunit gene *HTA1* (RTO4_12090) with the mRuby2 fluorescent protein at the native locus. The Hta1-mRuby2 tagged strains were generated using a combination of G418 resistance and a 5-fluorodeoxyuridine (FUDR) sensitive thymidine kinase counterselection marker flanked by 250 bp terminal repeat sequences. The Hta1-mRuby2 construct was transformed into the *CBR1-GFP* background using G418 to select for transformants. Transformed colonies were passaged one time on liquid yeast peptone dextrose (YPD, VWR) and plated onto yeast nitrogen base with $NH_4SO_4$ without amino acids (YNB, MP Biomedicals) + 1% glucose + 50 μg/mL FUDR (VWR) plates to select for loop-outs of the G418 and thymidine kinase markers. Colonies were screened by microscopy, PCR, and locus sequencing for correct integration of the mRuby2 tag and successful removal of the G418 and thymidine kinase markers. Since RTO4_16716 and RTO4_16717 are adjacent in the genome, the *bgl1*Δ *bgl2*Δ strain was generated via single transformation of the nourseothricin resistance cassette to remove both open reading frames from the genome, and the *bgl1*Δ *bgl2*Δ + *BGL1 BGL2* complement strain was generated from the *bgl1*Δ *bgl2*Δ strain by transforming a single genomic fragment containing both coding sequences. The *cbr1*Δ + $P_{AAC1}$-*BGL1* strain was generated via transformation of the *BGL1* open reading frame driven by the previously characterized constitutive *AAC1* (RTO4_12704) promoter [4] and the 35S rRNA terminator into the *YKU70* locus with a G418 resistance cassette for selection. All strains were confirmed by PCR. Complement, tagged, and constitutive expression strains were further confirmed by DNA sequencing of the transformed loci.

*R. toruloides growth experiments*

Colonies of *R. toruloides* grown on solid YPD were inoculated into liquid YPD and grown at 28°C with constant shaking at 250 rpm until exponential phase. Cells were pelleted at 4000 g and washed three times with YNB lacking a carbon source. Washed cells were used to inoculate 750 µL of media in 48-well plates at the following starting densities, as measured by an Eppendorf BioPhotometer with a 10 mm path length: 0.1 OD600/mL for initial carbon screening experiments (Fig S2) of wild type and *cbr1*Δ in YNB 1% acetate, YNB 1% arabinose, YNB 1% fructose, YNB 1% galacturonic acid, YNB 1% glucose, YNB 1% glutamate, YNB 1% glycine, YNB 1% L-alanine, YNB 1% maltose, YNB 1% sucrose, YNB 1% Tween-80, and YNB 1% xylose; 0.5 OD600/mL for the high density *cbr1*Δ + $P_{AAC1}$-*BGL1* growth experiment (Fig S6D); and 0.01 OD600/mL for all other growth experiments. Growth assays were incubated at 28°C with constant shaking (3 mm orbit, 600 rpm for experiments with manual, individual time points and 3 mm orbit, 425 rpm for kinetic growth experiments). OD600 measurements of manual, individual time points were taken with an Agilent BioTek Synergy HTX. Kinetic growth experiments were performed using an Agilent BioTek Epoch2 plate reader with OD600 measurements taken every 20 or 30 min depending on the experiment.

*N. crassa growth experiments*

For growth of *N. crassa* on trehalose and xylose with or without 2-deoxyglucose (TCI) wild type *mat A* (FGSC 2489) and Δ*clr-2 mat A* conidia were inoculated at $10^6$ conidia/mL into 250 mL flasks with 100 mL of filter sterilized Vogel's minimal medium [5] with 50 mM $NH_4Cl$ instead of 25 mM $NH_4NO_3$ (VMM $NH_4Cl$) with either 1% xylose or 1% trehalose. Filter sterilized 20% 2-deoxyglucose dissolved in water was added to the media to achieve 2-deoxyglucose concentrations of 0 μM, 119 μM, 238 μM, or 476 μM 2-deoxyglucose for media containing xylose as the sole carbon source and 2-deoxyglucose concentrations of 0 mM, 2.4 mM, 3.8 mM, or 7.6 mM for media containing trehalose as the sole carbon source. Cultures were grown at 28ºC with constant light and shaking at 200 rpm. After 24 h, biomass was autoclaved, harvested via filtration, washed with distilled water, dried for 24 h at 60°C in a drying oven, and weighed.

For growth of *N. crassa* on tricarboxylic acid (TCA) cycle intermediates and fucose, wild type *mat A* (FGSC 2489) and Δ*clr-2 mat A* conidia were inoculated at $10^6$ conidia/mL in 24-well deep-well round-bottomed plates with 3 mL of filter sterilized VMM $NH_4Cl$ with 1% succinate, 1% α-ketoglutarate, 1% malate, or 1% fucose. Cultures were incubated at 28ºC with constant light and shaking (200 rpm). Biomass from TCA cycle intermediates and fucose cultures was harvested by filtration 7 d and 10 d post inoculation, respectively. Biomass was washed with distilled water, dried for 24 h in a drying oven at 60ºC, and weighed.

*Microscopy*

Fluorescent microscopy was performed using a Nikon Eclipse Ti2-E inverted microscope equipped with a prime BSI express CMOS camera using a CFI Plan Apochromat 100X oil objective. *HTA1-mRuby2 CBR1-GFP* cells were grown to exponential phase in liquid YPD. Cells were pelleted at 4000 g and washed three times with YNB without a carbon source. Cells were inoculated into a 48-well plate with YNB + 1% w/v of the indicated carbon source at 0.2 OD600/mL, as measured by an Eppendorf BioPhotometer with a 10 mm path length, and incubated at 28ºC with shaking (3 mm orbit, 600 rpm) for 4 h. Aliquots of the cultures were placed onto agarose coated slides and imaged.

*RNA sequencing and transcript abundance*

Cells were grown to exponential phase in liquid YPD and washed three times in YNB without a carbon source. Two hundred fifty mL flasks containing 100 mL of YNB without a carbon source or YNB with the indicated carbon source were inoculated at 0.1 OD600/mL, as measured by an Eppendorf BioPhotometer with a 10 mm path length. Culture conditions and sequencing methods for each experiment are summarized in Table S4. Cultures were incubated at 28ºC with constant shaking (200 rpm) for the following times: 8 h for wild type and *cbr1*Δ cells exposed to YNB 1% cellobiose, YNB 1% succinate, YNB 1% fucose, and YNB without a carbon source (Fig 2), 4 h for wild type, *cbr1*Δ, and *bgl1*Δ *bgl2*Δ cells exposed to YNB 1% cellobiose (Fig 4), 24 h for wild type, *cbr1*Δ, and *tct1*Δ cells exposed to YNB 8.5 mM citrate (Fig 5), 24 h for wild type cells exposed to YNB 1% succinate (used to construct GFF file with improved 3' untranslated regions [UTR]), and 8 h for wild type cells exposed to VMM $NH_4Cl$ 2% glucose (used to construct GFF file with improved 3'UTRs). Cells were harvested using centrifugation in 50 mL conical tubes at 4000 g. The supernatants were removed, and the cell pellets were flash frozen in liquid nitrogen. Pellets were stored at -80ºC prior to RNA extraction. RNA was extracted using the Quick-RNA Fungal/Bacterial Miniprep Kit (Zymo) according to the manufacturer instructions, followed by either treatment with the TURBO DNA-free kit (Invitrogen) for standard RNA sequencing (RNAseq) or DNase I (NEB) treatment for 10 min at 37ºC and purification with the Monarch Spin RNA Cleanup Kit (NEB) for 3'RNAseq. RNA quality was assessed via agarose gel electrophoresis, and a subset of samples were also assessed for RNA quality using an Agilent BioAnalyzer at the Biotechnology Resource Center at Cornell University.

For wild type and *cbr1*Δ cells exposed to YNB 1% cellobiose, YNB 1% succinate, and YNB without a carbon source for 8h, RNA was submitted to the California Institute for Quantitative Biosciences at UC Berkeley (QB3-Berkeley) for library preparation and sequencing. Libraries were prepared using standard Illumina protocols and RNA was sequenced using an Illumina NextSeq 2000 with 150 bp paired end reads at a depth of approximately 20 million reads per sample. Raw reads were trimmed using fastp v. 0.23.4 with the --trim_poly_x setting [6]. The transcript abundance (transcripts per million, TPM) was quantified using Salmon v. 1.10.0 mapping to the *R. toruloides* IFO 0880 transcriptome (v4) [2] using the --validateMappings and --GCbias settings [7]. Differential expression was determined using DESeq2 v. 1.50.1 [8]. Genes were designated as differentially expressed if they had an average TPM greater than 1 in at least one of the two compared conditions, $p_{adj} < 0.05$, and $log_2$(fold change) greater than 2 or less than -2. These RNAseq data were deposited in the Gene Expression Omnibus (GEO) at the National Center for Biotechnology Information (NCBI) and are accessible through GEO series accession number GSE293943.

For wild type and *cbr1*Δ cells exposed to YNB 1% fucose; wild type, *cbr1*Δ, and *bgl1*Δ *bgl2*Δ cells exposed to YNB 1% cellobiose; wild type, *cbr1*Δ, and *tct1*Δ cells exposed to YNB 8.5 mM citrate; wild type cells exposed to YNB 1% succinate for 24 h; and wild type cells exposed to VMM $NH_4Cl$ 2% glucose for 8 h, RNA was submitted to the Biotechnology Resource Center at Cornell University for library preparation and sequencing. Libraries were prepared using the Lexogen QuantSeq 3' mRNA-Seq Library Prep Kit FWD V2, and RNA was sequenced using an Illumina NovaSeqX with 150 bp paired end reads at a depth of approximately 10 million reads per sample, except for wild type exposed to VMM $NH_4Cl$ 2% glucose for 8 h, which was sequenced on an Illumina NextSeq 500 with 75 bp single end reads at a depth of approximately 7.5 million reads per sample. Raw reads were trimmed using fastp v. 0.23.4 with the --trim_poly_x setting [6]. Reads were aligned to the *R. toruloides* IFO 0880 genome (v4) [2] using

STAR v. 2.7.11b [9] with --alignIntronMin set to 5, and BAM files were indexed using Samtools v. 1.20 [10].

Because the 3'UTRs were not perfectly annotated in the *R. toruloides* IFO 0880 genome (v4) [2], leading to undercounting 3'RNAseq reads, we used peaks2utr v. 1.4.1 [11] to annotate missing or trucated 3'UTRs. To improve 3'UTR annotation, we used all the wild type 3'RNAseq conditions (YNB 1% fucose, YNB 1% cellobiose, YNB 8.5 mM citrate, YNB 1% succinate [24 h], and VMM $NH_4Cl$ 2% glucose). All the BAM files output by STAR v. 2.7.11b were combined, and the resulting combined BAM file was used as the peaks2utr input. In total, 2,696 3'UTRs were annotated by peaks2utr and the resulting gff3 file is available in Dataset S7. GNU parallel v. 20170522 [12] was used to run HTSeq v. 2.0.9 [13] in parallel for read counting using the peaks2utr generated gff3 (Dataset S7) file with --nonunique set to random, -t set to mRNA, and -i set to ProteinId. Differential expression was determined using DESeq2 v. 1.50.1 [8]. Counts per million (CPM) values were determined by dividing the counts for a given gene by the sum of all counts for the sample and multiplying by $10^6$. Genes were designated as differentially expressed if they had an average CPM greater than 1 in at least one of the two compared conditions, $p_{adj} < 0.05$, and $\log_2$(fold change) great than 2 or less than -2. Euclidean distance and hierarchical clustering were performed on the $\log_2$-transformed CPM for all genes that had a $p_{adj} < 0.05$ in at least one pairwise comparison and an average CPM of at least 1 in one of the three strains for wild type, *cbr1*Δ, and *bgl1*Δ *bgl2*Δ exposed to cellobiose (Fig S8 and Dataset S3-S4). Expression data for wild type, *cbr1*Δ, and *tct1*Δ cells exposed to YNB 8.5 mM citrate was visualized using the ComplexHeatmap R package [14]. These RNAseq data were deposited in the GEO at NCBI and are accessible through GEO series accession number GSE313844.

*Functional enrichment analysis*

Kyoto Encyclopedia of Genes and Genomes (KEGG) [15] and Gene Ontology (GO) [16, 17] enrichment analyses were performed on genes differentially expressed between *cbr1*Δ and wild type cells exposed to cellobiose, succinate, carbon starvation, and fucose (Fig 2) using the enricher function of the clusterProfiler R package v. 4.18.1 with Benjamini-Hochberg multiple hypothesis testing correction and visualized using the clusterProfiler dotplot function [18]. GO annotations and protein sequences for the *R. toruloides* IFO 0880 genome (v4) [2] were downloaded from Mycocosm (https://mycocosm.jgi.doe.gov/Rhoto_IFO0880_4/Rhoto_IFO0880_4.home.html) [19]. KEGG Orthology IDs were assigned to protein sequences using BlastKOALA [20] and mapped to KEGG pathways [15] using the clusterProfiler gson_KEGG_mapper function [18]. Pathways that are not biologically relevant to fungi were manually removed (e.g., Insect hormone biosynthesis, Olfactory transduction, etc.). Genes were included in the functional enrichment analysis input if they were at least four-fold decreased in expression in the *cbr1*Δ strain relative to the wild type strain with $p_{adj} < 0.05$ and either an average TPM or CPM of at least 1 in at least one condition. BlastKOALA [20] assignments are available in Dataset S8.

*Reverse transcription quantitative PCR*

To measure *BGL1* expression in wild type cells during exposure to YNB 1% cellobiose, YNB 1% succinate, YNB 1% glucose, and YNB without a carbon source (Fig 3D), wild type cells were grown in YNB 1% glucose overnight, back-diluted into fresh YNB 1% glucose, and grown for 6-7 generations until reaching an OD600 ~1. Cells were then washed twice in YNB without a carbon source and inoculated at 0.8 OD600/mL, as measured by an Agilent BioTek Synergy HTX plate reader, in 100 mL YNB 1% cellobiose, YNB 1% succinate, YNB 1% glucose, or YNB without a carbon source in 250 mL flasks. Cultures were incubated at 28ºC with shaking at 200 rpm for 4 h. 300 µL of culture was then harvested, pelleted at 5500 g, supernatants were removed, and cells were flash frozen in liquid $N_2$ and stored at -80°C until RNA was extracted.

To measure *BGL1* expression in wild type cells during exposure to YNB lacking a carbon source, YNB 1% cellobiose, YNB 1% 2-deoxyglucose, YNB 100 μM 2-deoxyglucose, YNB 10 μM 2-deoxyglucose, YNB 1 μM 2-deoxyglucose, and YNB 0.1 μM 2-deoxyglucose (Fig 4D), exponentially growing cells were washed three times in YNB without a carbon source and 0.222 OD600/mL wild type cells, as measured by an Eppendorf BioPhotometer with a 10 mm path length, were inoculated in 90mL of YNB without a carbon source. Cultures were incubated at 28ºC with shaking at 200 rpm for 1 h. After 1 h, we added 10 mL of either YNB without a carbon source or YNB with 10X the final indicated concentrations of cellobiose or 2-deoxyglucose. Cultures were then incubated at 28ºC with shaking at 200 rpm. After 1 h, 10 mL of each culture was harvested, and after 4 h, 50 mL of each culture was harvested. Harvested cells were pelleted at 4000 g, the supernatants were removed, and cells were flash frozen in liquid $N_2$ and stored at -80ºC until RNA was extracted.

RNA was extracted using the Quick-RNA Fungal/Bacterial Miniprep Kit (Zymo) according to manufacturer instructions. Purified RNA was treated with DNase I (NEB) for 10 min at 37ºC. RNA was then purified using the Monarch® Spin RNA Cleanup Kit (NEB). RNA quality was assessed via agarose gel electrophoresis. RT-qPCR was performed using the Luna® Universal One-Step RT-qPCR Kit (NEB) according to manufacturer instructions with either 25 ng (carbon source experiment; Fig 3D) or 50 ng (2-deoxyglucose experiment; Fig 4D) RNA as template. Data were collected on a CFX96™ Real-Time System (Bio-Rad). Gene expression was normalized to *ACT1* (RTO4_14107) expression for all experiments. The following primer sets were used to quantify expression of each gene: *BGL1*, CGGGTGGAATCATGTGCTCGTA and CGTACCAGTCCGTGACGACAAAG; *ACT1*, CGTCCTCTCGCTCTATGCCTC and CTTGATGAGGTAGTCGGTCAGGTC.

*β-glucosidase enzyme activity assays*

Cells were grown to exponential phase in liquid YPD and washed three times in YNB without a carbon source. Washed cells were inoculated into 750 µL YNB 1% glucose, YNB 1% cellobiose, or YNB 1% succinate in 48-well plates at 0.8 OD600/mL, as measured by an Eppendorf BioPhotometer with a 10 mm path length. Wild type, *cbr1*Δ, *cbr1*Δ + *CBR1*, *bgl1*Δ *bgl2*Δ, *bgl1*Δ *bgl2*Δ + *BGL1 BGL2*, *bgl1*Δ, *bgl1*Δ + *BGL1*, and *bgl2*Δ cells were incubated for 20 h at 28°C with shaking (600 rpm, 3 mm orbit) (Fig 3C and Fig S6B-C). To measure the secreted β-glucosidase activity of *cbr1*Δ + $P_{AAC1}$-*BGL1* cells (Fig S6F), wild type and *cbr1*Δ + $P_{AAC1}$-*BGL1* cells were incubated at 28°C with shaking (600 rpm, 3 mm orbit) for 16 h to ensure both wild type and *cbr1*Δ + $P_{AAC1}$-*BGL1* cultures were in exponential phase at the time of the supernatant harvest. After 20 h or 16 h incubation, as described above, cultures were transferred to 1.5 mL Eppendorf tubes, and cells were pelleted by centrifuging at 14,000 g for 3 min. Supernatants were transferred to 0.2 µm centrifuge tube filters (CoStar Spin-X) and centrifuged at 14,000 g for 1 min. Filtered supernatants were stored at -20°C.

β-glucosidase activity was determined using a p-nitrophenyl-β-D-glucopyranoside (EMD Millipore Corp.) colorimetric enzyme activity assay. We added 50 µL of 100 mM sodium acetate (pH 5) and 25 µL of freshly prepared 20 mM p-nitrophenyl-β-D-glucopyranoside to each well of a 96-well plate on ice. We then added 25 µL of each supernatant to the wells of the 96-well plate and mixed by pipetting (Fig 3C and Fig S6F and S7A). To better determine whether β-glucosidase activity was present in *bgl1*Δ supernatants, we mixed 25 µL of 100 mM sodium acetate (pH 5) and 25 µL of freshly prepared 20 mM p-nitrophenyl-β-D-glucopyranoside dissolved in 100 mM sodium acetate (pH 5) and then added 50 µL of the indicated supernatant (Fig S6C). In each assay an equal number of mock reaction wells lacking supernatant were also prepared. Reactions were incubated at 37°C for 35 min followed by quenching with 100 µL of 200 mM $Na_2CO_3$. After quenching, 25 µL (Fig 3C and Fig S6F and S7A) or 50 µL (Fig S6C) of supernatant were added to the mock reactions as blanks for their respective reactions.

Absorbance was measured at 400 nm (OD400) using an Agilent BioTek Synergy HTX plate reader. The absorbance of the respective mock reaction was subtracted from each OD400, and OD400 values were normalized to the average OD400 of wild type cells grown on cellobiose for each experiment.

*Spent media growth assays*

Exponentially growing wild type cells were washed three times with YNB without a carbon source and inoculated at 0.01 OD600/mL, as measured by an Eppendorf BioPhotometer with a 10 mm path length, in 100 mL YNB 1% glucose, YNB 1% cellobiose, or YNB 1% succinate in 250 mL flasks. Cultures were grown to an OD600 of ~1, as measured by an Eppendorf BioPhotometer with a 10 mm path length, prior to centrifugation at 4000 g for 10 min. The supernatants were sterilized with vacuum bottle-top 0.22 μm filters (VWR). The β-glucosidase activity of each supernatant was assessed using the β-glucosidase assay described above. Filter sterilized supernatants were added to YNB lacking a carbon source or YNB 1% cellobiose to a final concentration of 6%, 12%, or 25% spent supernatant. Exponentially growing wild type, *cbr1*Δ, and *bgl1*Δ *bgl2*Δ cells were washed three times with YNB without a carbon source and inoculated at 0.01 OD600/mL, as measured by an Eppendorf BioPhotometer with a 10 mm path length, in 750 μL of media containing the indicated concentrations of filtered supernatant in 48-well plates. Cultures were incubated at 28ºC with shaking at 600 rpm (3 mm orbit), and OD600 measurements were taken at the indicated times.

### SI FIGURES

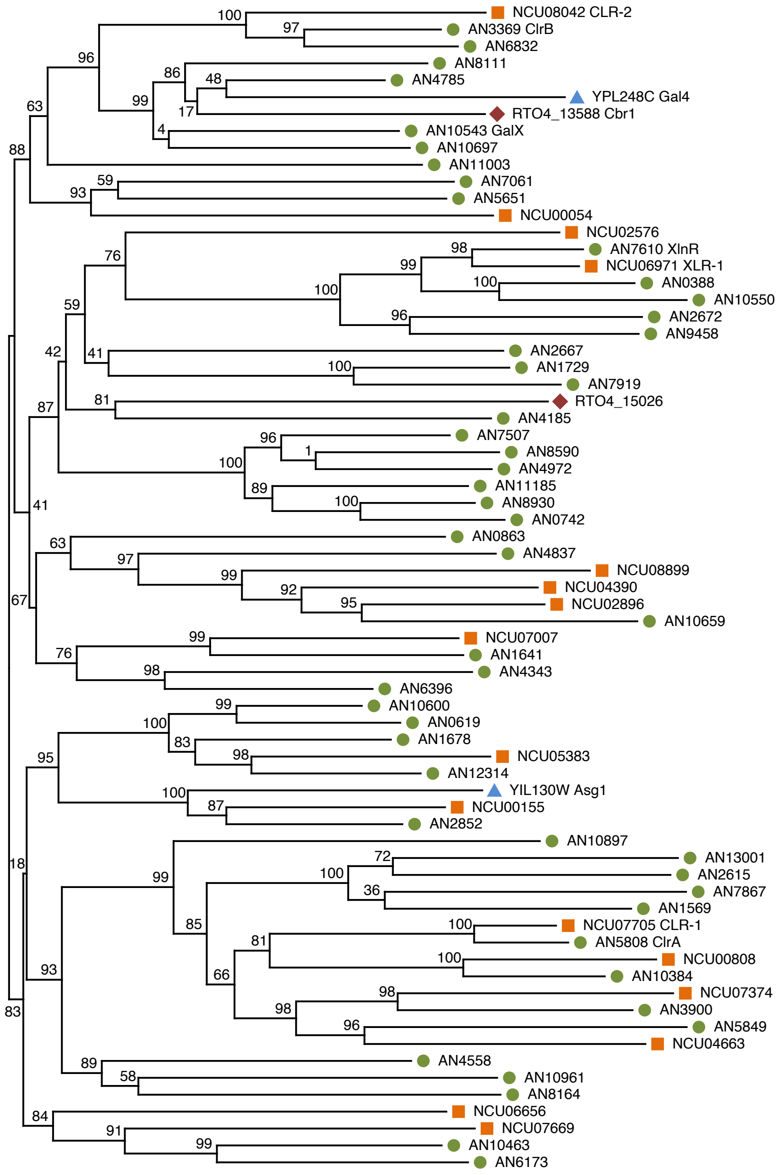


**Fig S1. Phylogenetic tree of homologs of transcription factors regulating cellulose and hemicellulose utilization.** Phylogenetic tree of CLR-1/ClrA, CLR-2/ClrB, and XLR-1/XlnR homologs in *Rhodotorula (Rhodosporidium) toruloides*, *Neurospora crassa*, *Aspergillus nidulans*, and *Saccharomyces cerevisiae* built using the maximum likelihood method based on the Jones Taylor-Thornton matrix-based model using FastTree [21]. Numbers at nodes indicate the reliability of each split in the tree computed using the Shimodaira-Hasegawa test on three alternate topologies around that split with 1,000 resamples. *R. toruloides* proteins are indicated with maroon diamonds. *N. crassa* proteins are indicated with orange squares. *A. nidulans* proteins are indicated with green circles. *S. cerevisiae* proteins are indicated with blue triangles.

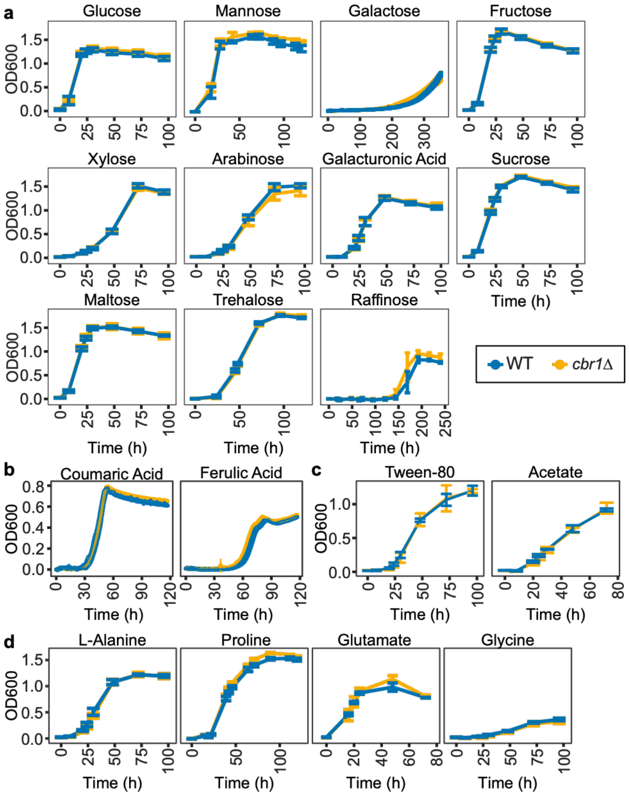


**Fig S2. *CBR1* (RTO4_13588) is not required for growth on most carbon sources.** Growth (OD600) of wild type (WT) and *cbr1*Δ cells grown in YNB with the indicated carbon source. All carbon sources were added at 1% w/v unless indicated otherwise: (**A**) sugars, (**B**) 0.2% w/v phenolic lignin building blocks, (**C**) carbon sources metabolized through acetyl-CoA, and (**D**) amino acids. Lines are the average and bars or colored bands are the standard deviation of three biological replicates.

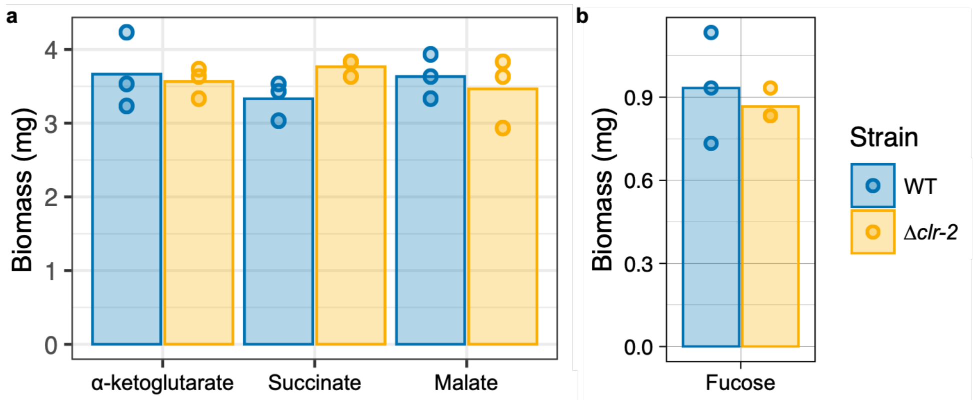


**Fig S3. *N. crassa clr-2* is not required for utilization of TCA cycle intermediates or fucose.** Mycelial dry weight of wild type (WT) and Δ*clr-2 N. crassa* cells directly inoculated into 3 mL of (**A**) VMM $NH_4Cl$ 1% TCA cycle intermediates grown for 7 d post inoculation or (**B**) VMM $NH_4Cl$ 1% fucose grown for 10 d post inoculation. Bars are the average and dots are the individual data points of three biological replicates.

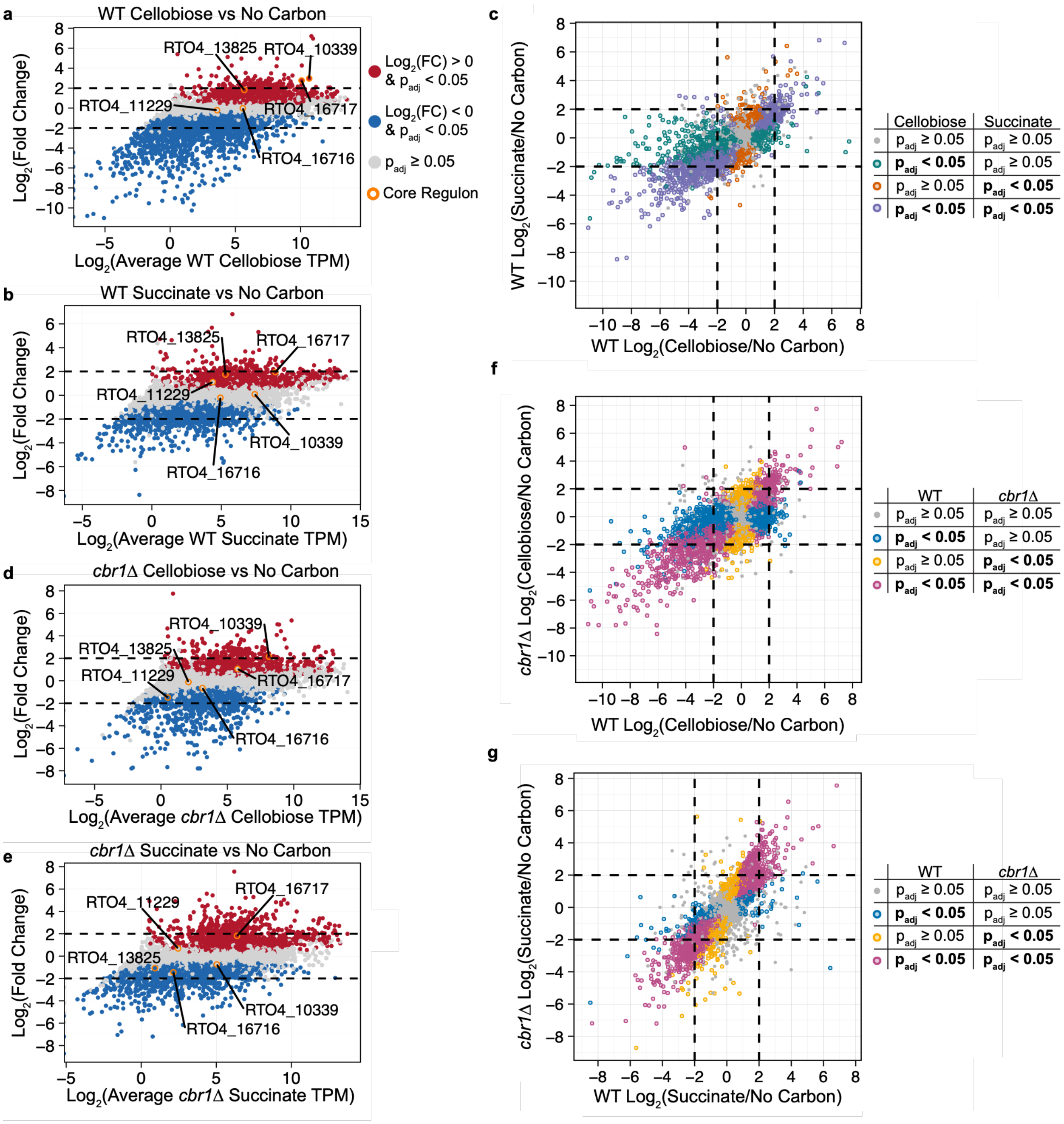


**Fig S4. Exposure to cellobiose and succinate yield a similar transcriptional profile that is largely independent of *CBR1*.** (**A-B**) Scatterplots of the $\log_2$(average gene expression) during exposure to (**A**) cellobiose or (**B**) succinate vs $\log_2$(fold change) in gene expression during exposure to (**A**) cellobiose or (**B**) succinate compared to carbon starvation in wild type cells. (**C**) Scatterplot of the $\log_2$(fold change) in gene expression in wild type cells exposed to cellobiose compared to carbon starvation vs succinate compared to carbon starvation. Genes with a $p_{adj}$ <

0.05 in succinate only are colored orange, a $p_{adj} < 0.05$ in cellobiose only are colored green, and a $p_{adj} < 0.05$ in both cellobiose and succinate are colored purple. Genes with a $p_{adj} \geq 0.05$ in all comparisons are colored grey. (**D-E**) Scatterplots of the $\log_2$(average gene expression) during exposure to (**D**) cellobiose or (**E**) succinate vs $\log_2$(fold change) in gene expression during exposure to (**D**) cellobiose or (**E**) succinate compared to carbon starvation in *cbr1*Δ cells. (**F-G**) Scatterplots of the $\log_2$(fold change) in gene expression in wild type vs *cbr1*Δ cells exposed to (**F**) cellobiose or (**G**) succinate compared to carbon starvation. Genes with a $p_{adj} < 0.05$ in wild type cells only are colored blue, a $p_{adj} < 0.05$ in *cbr1*Δ only are colored gold, and a $p_{adj} < 0.05$ in both *cbr1*Δ and wild type are colored magenta. Genes with a $p_{adj} \geq 0.05$ in all comparisons are colored grey. For (**A**), (**B**), (**D**), and (**E**), genes with a $p_{adj} < 0.05$ are colored red if they had higher expression in the indicated carbon source compared to carbon starvation and blue if they had lower expression in the indicated carbon source compared to carbon starvation. The five genes determined to be activated by *CBR1* across multiple carbon sources (*CBR1* core regulon) are highlighted in orange and labeled with their respective protein IDs. FC stands for fold change and TPM stands for transcripts per million. RNAseq experiments had three biological replicates.

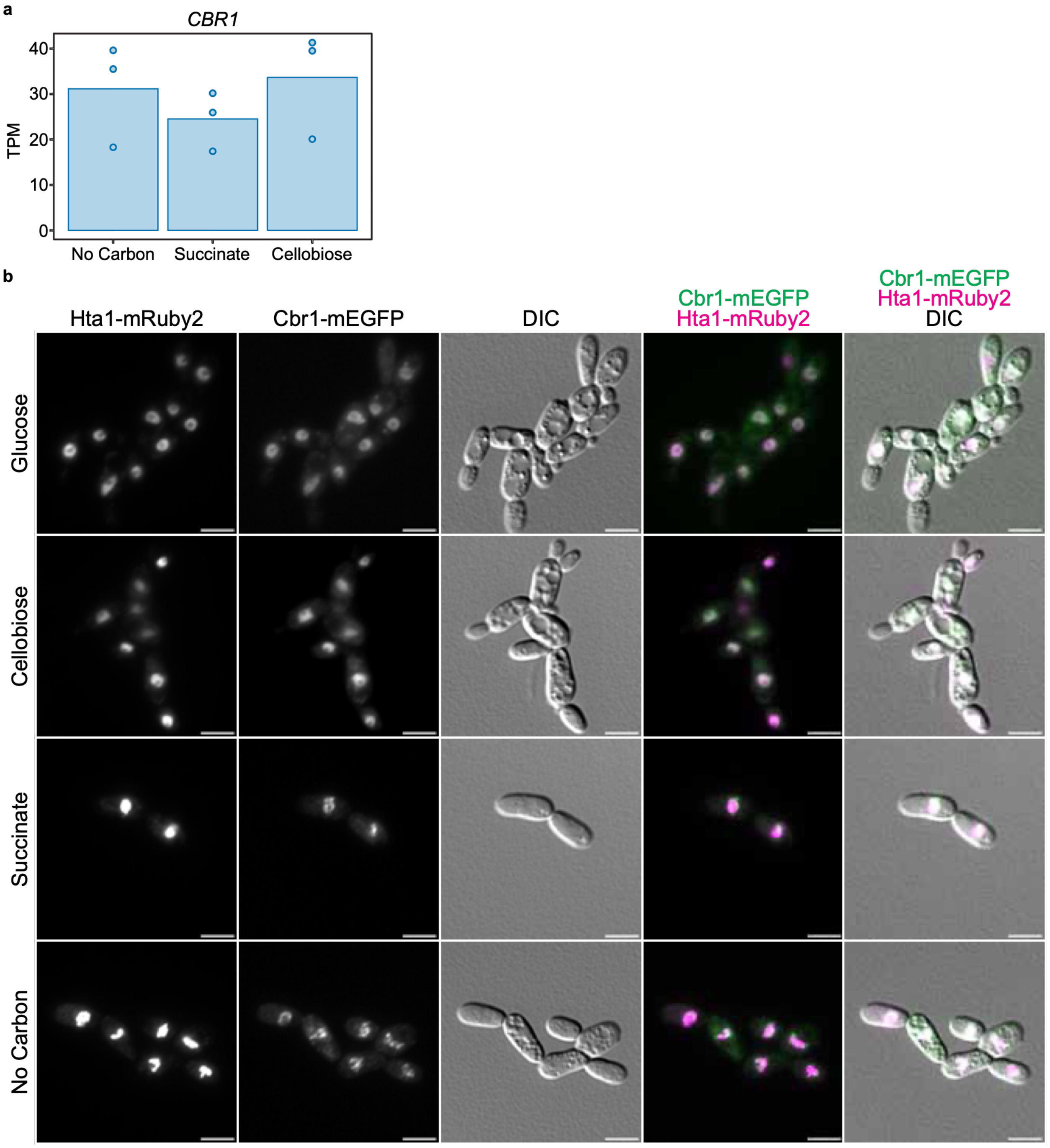


**Fig S5. Cbr1 is present in the nucleus during exposure to glucose, cellobiose, succinate, and carbon starvation.** (**A**) Expression (transcripts per million, TPM) of *CBR1* during exposure to media containing succinate, cellobiose, or lacking a carbon source (no carbon). Bars are the average and dots are the individual data points of three biological replicates. (**B**) Cells with the histone subunit Hta1 (H2A) tagged with mRuby2 (Hta1-mRuby2) and Cbr1 tagged with GFP

(Cbr1-mEGFP) were exposed to the indicated carbon source for 4 h prior to imaging. Fluorescent microscopy was performed using a Nikon Eclipse Ti2-E inverted microscope equipped with a prime BSI express CMOS camera using a CFI Plan Apochromat 100X oil objective. The scale bar is 5 μm. Images are representative of three biological replicates.

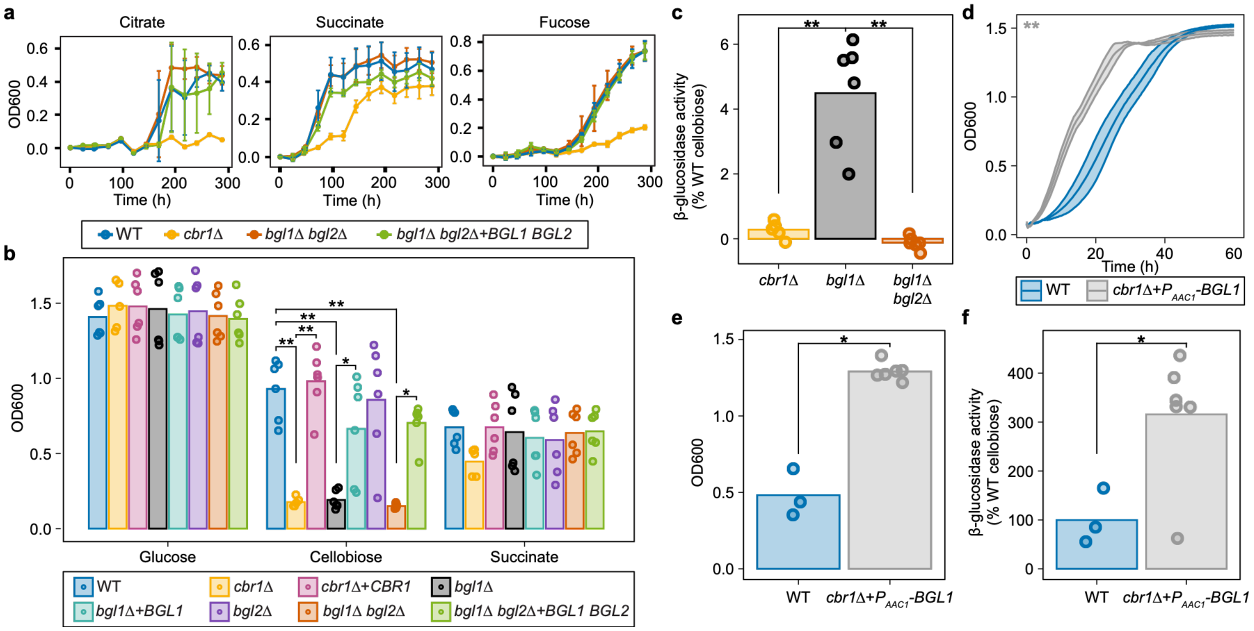


**Fig S6. Cbr1-dependent expression of secreted β-glucosidase genes *BGL1* (RTO4_16717) and *BGL2* (RTO4_16716) is necessary for cellobiose but not TCA cycle intermediate utilization.** (**A**) Growth curves (OD600) of wild type, *cbr1*Δ, *bgl1*Δ *bgl2*Δ, and *bgl1*Δ *bgl2*Δ + *BGL1 BGL2* cells grown on YNB 8.5 mM citrate, YNB 1% succinate, or YNB 1% fucose. (**B**) Cell density (OD600) at the time of supernatant harvest (20 h) for cultures corresponding to Fig 3C and Fig S6C β-glucosidase activity. (**C**) β-glucosidase activity of supernatants from *cbr1*Δ, *bgl1*Δ, and *bgl1*Δ *bgl2*Δ strains grown on cellobiose, normalized to the β-glucosidase activity present in the culture supernatant of wild type cells grown on cellobiose. Twice as much supernatant was used as input for these β-glucosidase activity assays than for all other β-glucosidase activity assays (see Text S1). (**D**) Growth curve (OD600) of wild type and *cbr1*Δ + $P_{AAC1}$-*BGL1* cells on YNB 1% cellobiose with a starting inoculum 50-fold higher than in Fig 3B. (**E**) Cell density (OD600) of wild type and *cbr1*Δ + $P_{AAC1}$-*BGL1* cells grown in YNB 1% cellobiose at the time of the supernatant harvest (16 h) for cultures corresponding to the β-glucosidase activity in culture supernatants in Fig S6F. (**F**) β-glucosidase activity present in the culture supernatant of wild type and *cbr1*Δ + $P_{AAC1}$-*BGL1* cells grown on YNB 1% cellobiose for 16 h. (**A**

and **D**) Lines are the average and bars or colored bands are the standard deviation of three biological replicates. (**B**, **C**, **E**, and **F**) Bars are the average and points are the individual data points of (**B-C**) five biological replicates for *cbr1*Δ cells and six biological replicates for all other strains or (**E-F**) three biological replicates for wild type cells and six biological replicates for *cbr1*Δ + $P_{AAC1}$-*BGL1* cells. *$p_{adj} < 0.05$ and **$p_{adj} < 10^{-4}$ as determined by (**A** and **D**) pairwise comparisons of estimated marginal means with a holm multiple comparison correction, (**B**) two-way ANOVA with a TukeyHSD post-hoc test, (**C**) one-way ANOVA with a TukeyHSD post-hoc test, or (**E** and **F**) Welch's two-sample *t*-test.

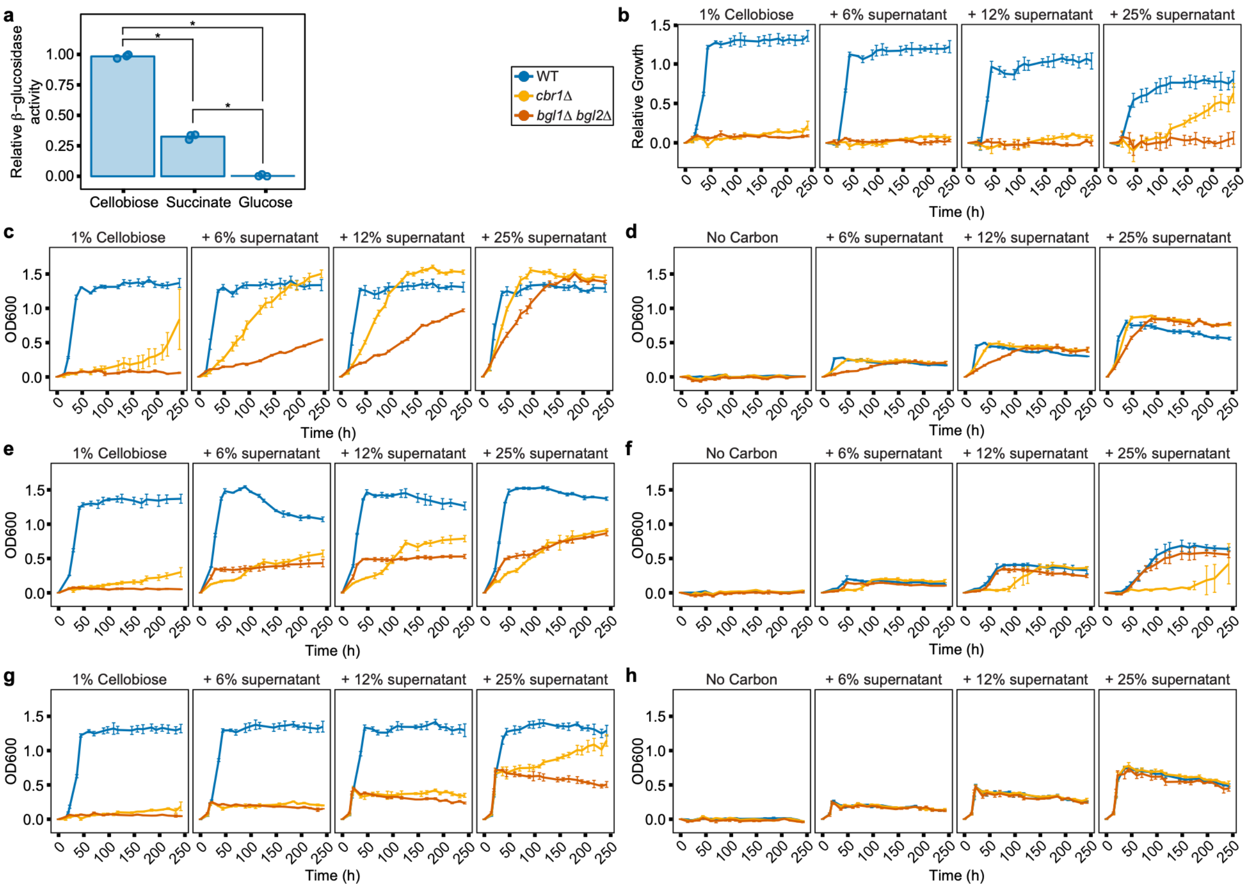


**Fig S7. Secreted β-glucosidases in spent supernatants are sufficient to complement the growth of *cbr1*Δ and *bgl1*Δ *bgl2*Δ cells on cellobiose.** (**A**) β-glucosidase activity of culture supernatants of wild type cells grown in YNB with 1% of the indicated carbon source. (Supernatants used to supplement media in Fig 3E, 3F, and Fig S6B-S6H.) *$p_{adj} < 10^{-6}$ as determined by a one-way ANOVA with a TukeyHSD post-hoc test. (**B**) Normalized growth (OD600) of the indicated strains in YNB 1% cellobiose supplemented with the indicated concentration of spent supernatant from wild type (WT) cells grown in YNB 1% glucose. To account for growth due to nutrients remaining in the supplemented supernatant, growth was normalized by subtracting the growth of each replicate on media lacking a carbon source supplemented with the same concentration of spent supernatant. (**C, E,** and **G**) Growth (OD600) of wild type, *cbr1*Δ, and *bgl1*Δ *bgl2*Δ cells grown in YNB 1% cellobiose supplemented with the indicated concentration of culture supernatant from wild type cells grown on (**C**) cellobiose, (**E**)

succinate, or (**G**) glucose. (**D, F,** and **H**) Growth (OD600) of wild type, *cbr1*$\Delta$, and *bgl1*$\Delta$ *bgl2*$\Delta$ cells grown in YNB lacking a carbon source supplemented with the indicated concentration of culture supernatant from wild type cells grown on (**D**) cellobiose, (**F**) succinate, or (**H**) glucose. (**A**) Bars are the average and points are the individual data points of three biological replicates. (**B-H**) Lines are the average and bars are the standard deviation of three biological replicates.

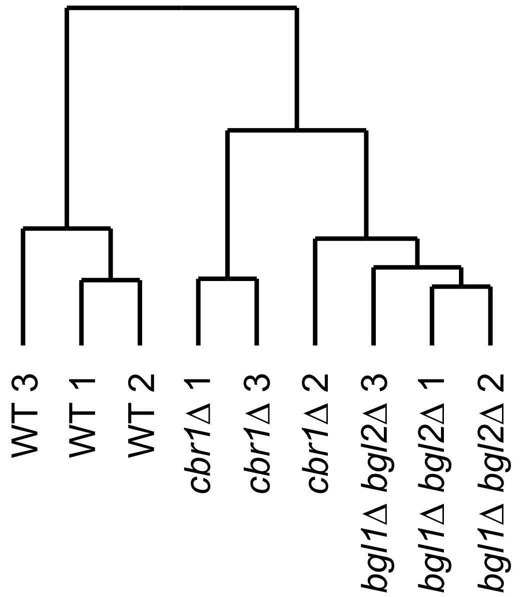


**Fig S8. The transcriptional profile of cells lacking *BGL1* and *BGL2* is more similar to the transcriptional profile of *cbr1*Δ cells than wild type cells during exposure to cellobiose.** Hierarchical clustering using Euclidean distance of gene expression data of wild type, *cbr1*Δ, and *bgl1*Δ *bgl2*Δ cells exposed to cellobiose for 4 h for all genes with $p_{adj} < 0.05$ in at least one of the following pairwise comparisons: wild type cells compared to *bgl1*Δ *bgl2*Δ cells, wild type cells compared to *cbr1*Δ cells, or *bgl1*Δ *bgl2*Δ cells compared to *cbr1*Δ cells.

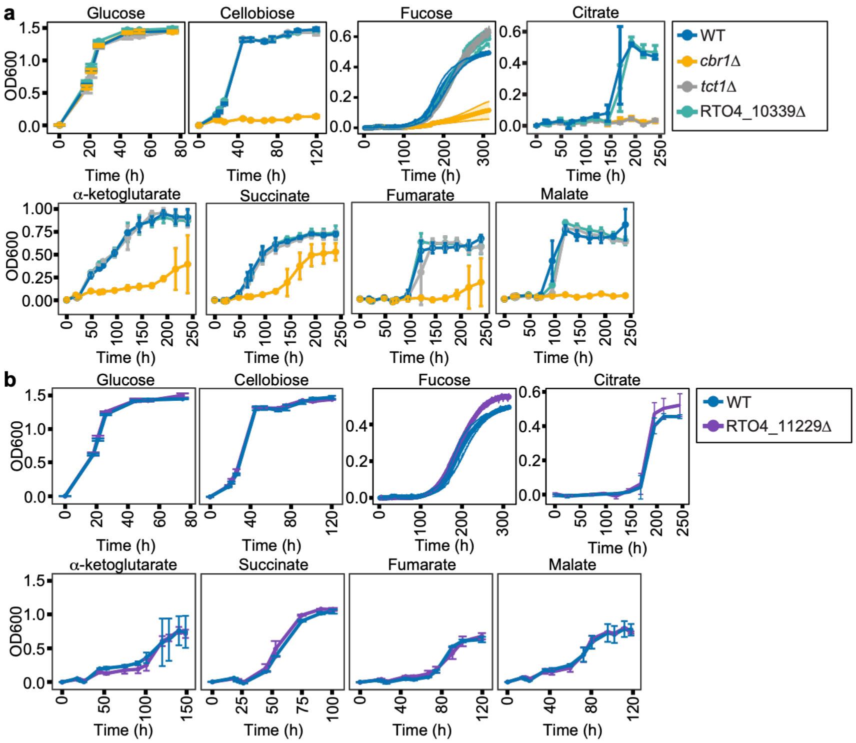


**Fig S9. Growth of *tct1*Δ (RTO4_13825Δ), RTO4_10339Δ, and RTO4_11229Δ cells on TCA cycle intermediates, cellobiose, and fucose.** (**A** and **B**) Growth (OD600) of the indicated strains in YNB with 1% of the indicated carbon source for all carbon sources except citrate, which is at 8.5 mM. Lines are the average and bars or colored bands are the standard deviation of three biological replicates.

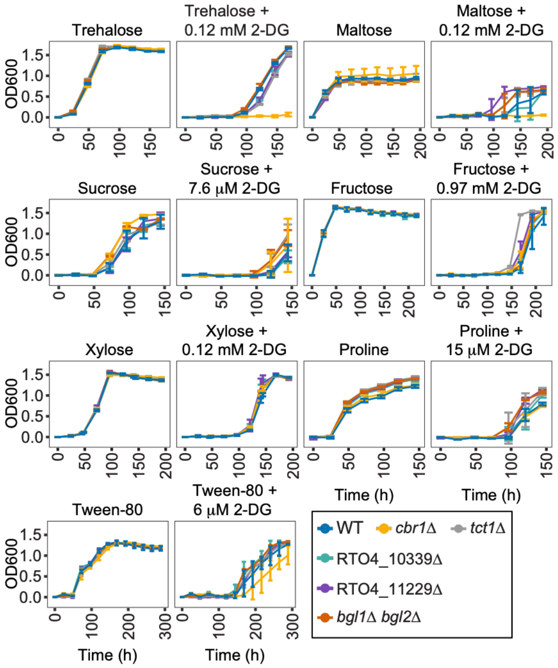


**Fig S10. Deletions of genes in the *CBR1* core regulon do not result in a growth phenotype when 2-deoxyglucose is added to media containing nonpreferred carbon sources.** Growth (OD600) of the indicated strains in YNB with 1% of the indicated carbon source supplemented with the indicated concentration of 2-deoxyglucose. Lines are the average and bars are the standard deviation of three biological replicates.

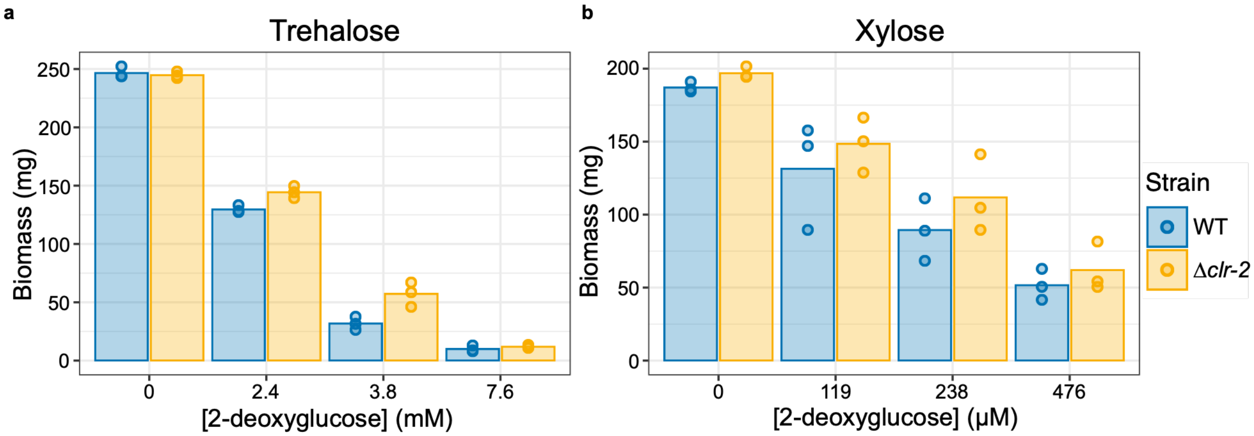


**Fig S11. *N. crassa* CLR-2 does not regulate carbon catabolite repression in *N. crassa*.** Mycelial dry weights of wild type and Δ*clr-2* *N. crassa* cells directly inoculated into 100 mL VMM $NH_4Cl$ containing 1% (**A**) trehalose or (**B**) xylose supplemented with the indicated concentration of 2-deoxyglucose and grown for 24 h prior to harvesting. Bars are the averages and dots are the individual data points of three biological replicates.

### SI TABLES

**Table S1. *R. toruloides* strains used in this study.**

| Strain name | Genotype | Source |
|---|---|---|
| Wild type | *yku70*(RTO4_11920)Δ::$Hyg^R$* | [1] |
| *cbr1*Δ | *cbr1*(RTO4_13588)Δ::$Nat^R$**<br>*yku70*Δ::$Hyg^R$ | This study |
| *cbr1*Δ + *CBR1* | *cbr1*Δ::$Nat^R$<br>*yku70*Δ::*CBR1*:$G418^R$*** | This study |
| *HTA1-mRuby2 CBR1-mEGFP* | *HTA1*(RTO4_12090)-*mRuby2*<br>*CBR1-mEGFP*:$Nat^R$<br>*yku70*Δ::$Hyg^R$ | This study |
| RTO4_10339Δ | RTO4_10339Δ::$G418^R$<br>*yku70*Δ::$Hyg^R$ | This study |
| RTO4_11229Δ | RTO4_11229Δ::$G418^R$<br>*yku70*Δ::$Hyg^R$ | This study |
| *tct1*Δ | *tct1*(RTO4_13825)Δ::$G418^R$<br>*yku70*Δ::$Hyg^R$ | This study |
| *tct1*Δ + *TCT1* | *tct1*Δ::$G418^R$<br>*yku70*Δ::*TCT1*:$Nat^R$ | This study |
| *bgl1*Δ | *bgl1*(RTO4_16717)Δ::$Nat^R$<br>*yku70*Δ::$Hyg^R$ | This study |
| *bgl1*Δ + *BGL1* | *bgl1*Δ::$Nat^R$<br>*yku70*Δ::*BGL1*:$G418^R$ | This study |
| *bgl2*Δ | *bgl2*(RTO4_16716)Δ::$Nat^R$<br>*yku70*Δ::$Hyg^R$ | This study |
| *bgl1*Δ *bgl2*Δ | *bgl1*Δ *bgl2*Δ::$Nat^R$****<br>*yku70*Δ::$Hyg^R$ | This study |
| *bgl1*Δ *bgl2*Δ + *BGL1 BGL2* | *bgl1*Δ *bgl2*Δ::$Nat^R$****<br>*yku70*Δ::*BGL1 BGL2*:$G418^R$**** | This study |
| *cbr1*Δ + $P_{AAC1}$-*BGL1* | *cbr1*Δ::*NatR*<br>*yku70*Δ::$P_{AAC1}$-*BGL1*:$G418^R$ | This study |

*$Hyg^R$ stands for hygromycin resistance cassette.

**$Nat^R$ stands for nourseothricin resistance cassette.

***$G418^R$ stands for G418 resistance cassette.

****RTO4_16716 and RTO4_16717 are adjacent in the genome, so the *bgl1*Δ *bgl2*Δ strain was generated via single transformation of the nourseothricin resistance cassette to remove both open reading frames from the genome, and a genomic fragment containing both genes was used to complement the deletion of *BGL1* and *BGL2* in the *bgl1*Δ *bgl2*Δ + *BGL1 BGL2* strain.

**Table S2. *N. crassa* strains used in this study.**

| **Strain name** | **Genotype** | **Source** |
|---|---|---|
| Wild type | Wild type *mat A* | FGSC* 2489 [22] |
| $\Delta$*clr-2* | $\Delta$*clr-2* (NCU08042)::$Hyg^R$ *mat A* | FGSC 15834 [23] |

*FGSC stands for Fungal Genetics Stock Center (3).

**Table S3. Carbon sources used in this study.**

| **Carbon source** | **Company** |
|---|---|
| 2-deoxyglucose | TCI |
| Acetate | EMD |
| Alanine | BeanTown Chemical |
| Arabinose | Chem-Impex |
| Cellobiose | RPI |
| Citrate | VWR |
| Coumaric acid | TCI |
| Ferulic acid | Ambeed |
| Fructose | VWR |
| Fucose | TCI |
| Fumaric acid | BeanTown Chemical |
| Galactose | BeanTown Chemical |
| Galacturonic acid | Thermo |
| Glucose | VWR |
| Glutamate | Sigma |
| Glycine | EM Science |
| Malic acid | TCI |
| Maltose | BeanTown Chemical |
| Mannose | BeanTown Chemical |
| Proline | BeanTown Chemical |
| Succinic acid | VWR |
| Sucrose | Fisher |
| Trehalose | TCI |
| Tween-80 (Polysorbate 80) | VWR |
| Xylose | Thermo |
| $\alpha$-Ketoglutarate | Alfa Aesar |

**Table S4. RNA sequencing method.**

| Media | Strains | Hours exposed to carbon source | RNA sequencing method | Relevant figure |
|---|---|---|---|---|
| YNB 1% cellobiose, YNB 1% succinate, and YNB no carbon | Wildtype and *cbr1Δ* | 8 h | Whole transcript mRNA sequencing | Fig 2A-2C, 2E, S4, S5A, and Dataset S1 and S2 |
| YNB 1% fucose | Wildtype and *cbr1Δ* | 8 h | 3'mRNA sequencing | Fig 2D, 2E, and Dataset S1 and S2 |
| YNB 1% cellobiose | Wildtype, *cbr1Δ*, and *bgl1Δ bgl2Δ* | 4 h | 3'mRNA sequencing | Fig 4A-4C and Dataset S3 and S4 |
| YNB 8.5 mM citrate | Wildtype, *cbr1Δ,* and *tct1Δ* | 24 h | 3'mRNA sequencing | Fig 5B and Dataset S5 and S6 |
| YNB 1% succinate | Wild type | 24 h | 3'mRNA sequencing | Only used to annotate 3'UTRs |
| VMM $NH_4Cl$ 2% glucose | Wild type | 8 h | 3'mRNA sequencing | Only used to annotate 3'UTRs |

**SI DATASETS**

**Dataset S1. Expression of all genes during exposure of wild type and *cbr1*Δ cells to cellobiose, succinate, and media lacking a carbon source for 8 h as measured by standard RNAseq and wild type and *cbr1*Δ cells exposed to fucose for 8 h as measured by 3'RNAseq.**

**Dataset S2. Differential gene expression calculated by DESeq2 [8] in the indicated comparisons as measured by standard RNAseq of wild type and *cbr1*Δ cells exposed to cellobiose, succinate, and media lacking a carbon source for 8 h or 3'RNAseq of wild type and *cbr1*Δ cells exposed to fucose for 8 h.**

**Dataset S3. Expression of all genes during exposure of wild type, *cbr1*Δ, and *bgl1*Δ *bgl2*Δ cells to cellobiose for 4 h as measured by 3'RNAseq.**

**Dataset S4. Differential gene expression calculated by DESeq2 [8] in the indicated comparisons as measured by 3'RNAseq of wild type, *cbr1*Δ, and *bgl1*Δ *bgl2*Δ cells exposed to cellobiose for 4 h.**

**Dataset S5. Expression of all genes during exposure of wild type, *cbr1*Δ, and *tct1*Δ cells to citrate for 24 h as measured by 3'RNAseq.**

**Dataset S6. Differential gene expression calculated by DESeq2 [8] in the indicated comparisons as measured by 3'RNAseq of wild type, *cbr1*Δ, and *tct1*Δ cells exposed to citrate for 24 h.**

**Dataset S7. GFF3 file made using peaks2utr [11] with improved 3'UTR annotations generated using the 3'RNAseq data of wild type cells exposed to: YNB 1% fucose, YNB 1% cellobiose, YNB 8.5 mM citate, YNB 1% succinate, and VMM $NH_4Cl$ 2% glucose. We subsequently used this GFF3 file to analyze all 3'RNAseq data.**

**Dataset S8. KEGG orthology IDs for genes in the *R. toruloides* genome assigned by BlastKOALA [20].**

**Dataset S9. Numerical values to generate all bar and line graphs.**

23. Colot HV, Park G, Turner GE, Ringelberg C, Crew CM, Litvinkova L, et al. A high-throughput gene knockout procedure for *Neurospora* reveals functions for multiple transcription

factors. Proc Natl Acad Sci U S A. 2006;103(27):10352-7. Epub 2006/06/28. doi: 10.1073/pnas.0601456103. PubMed PMID: 16801547; PubMed Central PMCID: PMC1482798.
